# Precision Medicine Advancements Using Whole Genome Sequencing, Noninvasive Whole Body Imaging, and Functional Diagnostics

**DOI:** 10.1101/497560

**Authors:** Ying-Chen Claire Hou, Hung-Chun Yu, Rick Martin, Natalie M. Schenker-Ahmed, Michael Hicks, Elizabeth T. Cirulli, Isaac V. Cohen, Thomas J. Jönsson, Robyn Heister, Lori Napier, Christine Leon Swisher, Saints Dominguez, Haibao Tang, Weizhong Li, Jaime Barea, Christina Rybak, Emily Smith, Keegan Duchicela, Michael Doney, Pamila Brar, Nathaniel Hernandez, Ewen F. Kirkness, Andrew M. Kahn, J. Craig Venter, David S. Karow, C. Thomas Caskey

## Abstract

We report the results of a three-year precision medicine study that enrolled 1190 presumed healthy participants at a single research clinic. To enable a better assessment of disease risk and improve diagnosis, a precision health platform that integrates non-invasive functional measurements and clinical tests combined with whole genome sequencing (WGS) was developed. The platform included WGS, comprehensive quantitative non-contrast whole body (WB) and brain magnetic resonance imaging/angiography (MRI/MRA), computed tomography (CT) coronary artery calcium scoring, electrocardiogram, echocardiogram, continuous cardiac monitoring, clinical laboratory tests, and metabolomics. In our cohort, 24.3% had medically significant genetic findings (MSF) which may contribute to increased risk of disease. A total of 206 unique medically significant variants in 111 genes were identified, and forty individuals (3.4%) had more than one MSF. Phenotypic testing revealed: 34.2% of our cohort had a metabolomics profile suggestive of insulin resistance, 29.2% had elevated liver fat identified by MRI, 16.4% had clinically important cardiac structure or cardiac function abnormalities on cardiac MRI or ECHO, 8.8% had a high cardiovascular risk on CT coronary artery calcium scoring (Agatston calcium score > 400, Relative Risk of 7.2), 8.0% had arrhythmia found on continuous rhythm monitoring, 6.5% had cardiac conduction disorders found on EKG, 2% had previously undetected tumors detected by WB MRI, and 2.5% had previously undetected aneurysms detected by non-contrast MRI/MRA. Using family histories, personal histories, and test results, clinical and phenotypic findings were correlated with genomic findings in 130 study participants (63.1%) with high to moderate penetrance variants, suggesting the precision health platform improves the diagnostic process in asymptomatic individuals who were at risk. Cardiovascular and endocrine diseases achieved considerable clinical associations between MSFs and clinical phenotypes (89% and 72%, respectively). These findings demonstrate the value of integrating WGS and noninvasive clinical assessments for a rapid and integrated point-of-care clinical diagnosis of age-related diseases that contribute to premature mortality.

## INTRODUCTION

The completion of the Human Genome Project provided a wealth of genetic information, allowing an individual’s genetic variability to be considered for the purpose of precision medicine^1,2^. The goals of precision medicine include the improvement in prediction, prevention, diagnosis, and treatment of diseases^3^. Several ongoing studies, including the US Precision Medicine Initiative (All of Us)^3^, the Electronic Medical Records and Genomics (eMERGE) Network^4^, the Million Veteran Program^5^, the Kaiser Permanente Research Program on Genes, Environment and Health (RPGEH)^6^, the UK Biobank Initiative^7^, and the DiscovEHR collaboration^8^, are investigating the impact of integrating genomics data and clinical information to improve health and prevent disease. The initial insights from these precision medicine initiatives have been shared with the scientific community. For example, the initial results from DiscovEHR study have shown that 3.5% of individuals in the cohort had clinically actionable genetic variants in 76 genes, and the detection of pathogenic variation identifies at-risk patients who can benefit from proactive treatments^8^. DiscovEHR findings also showed that ~65% of individuals who carry a pathogenic variant had associated phenotypes observed in their medical records^8^. The initial insights into the delivery of genomic knowledge and approaches for clinical decision and clinical care also have been discussed^9, 10, 11, 12^.

Historically, the use of a retrospective observational cohort study design was the initial approach to study genomic and health associations. The phenotypic data of this approach may be limited to health outcomes and medical data available to a specific disease or trait; thus, findings may need to be focused on case-control analyses^13, 14^. Recent approaches of using longitudinal electronic health records have allowed the assessment of genetic variation in a wide range of diseases and the identification of loss of function variants in humans that improve our understanding of previously undiscovered biological functions and the development of therapeutic targets^4, 8, 7^. Molecular technologies, including metabolomics (metabolites), transcriptomics (RNA), proteomics (proteins), and epigenomics also have been employed to reveal the functional significance of genomic variations^15, 16, 17, 18^. In particular, the integration of DNA sequencing with metabolomics has proven useful for discoveries of disease-associated genes and biomarkers^19, 20, 21^.

Advancements in imaging technologies allow new approaches for detecting, measuring and analyzing a wide range of health information, including quantitative and qualitative analysis of the brain, neurological, cardiovascular, endocrine, liver disease, body composition, and tumor detection^22, 23, 24, 25, 26, 27^. Oncology has been a major beneficiary of early cancer detection and therapy management^22, 28^. Non-invasive and targeted imaging also has been instrumental for the early detection of neurological and cardiovascular disorders^26,28,29^. Whole body and brain magnetic resonance imaging (MRI) have been shown to derive quantitative imaging biomarkers^30^ which can be integrated with genomic variation and blood biomarkers for disease risk prediction and can be used to assess an individual’s health status. For example, volumes of brain regions derived from segmentation of an individual’s brain MRI can be used as quantitative biomarkers to assess an individual’s brain health^31^. Segmentation of muscle and adipose tissues into compartments such as lean muscle mass, visceral adipose tissue, and subcutaneous adipose tissue, can be used as biomarkers to assess an individual’s metabolic health, including risk for Type 2 diabetes^32, 33^. These quantitative and qualitative imaging biomarkers promise to yield significant benefits as a tool for precision medicine.

In this study, our efforts were focused on allowing better assessment of disease risk, better understanding of disease mechanisms, and individualized optimal therapy for the disease, which are the goals of the precision medicine initiative. Thus, in addition to WGS that has been previously incorporated into precision medicine efforts, we employed advanced imaging, omics technologies, and clinical tests to expand upon previous efforts. The advanced technologies included whole body and brain magnetic resonance imaging (MRI), computed tomography (CT) scan, electrocardiogram, echocardiogram, continuous cardiac monitoring, clinical laboratory tests, metabolomics, and microbiome testing, providing an unprecedented depth of data for each individual. In this paper, we combine WGS with deep quantitative phenotypic testing for early detection of age-related chronic diseases associated with premature mortality, including cancer, cardiovascular, endocrine, cirrhosis, and neurological disorders. Additionally, we share our initial findings on personalized imaging biomarkers and drug response.

## MATERIALS AND METHODS

### Study Population

We enrolled active adults ≥18 yrs old (without acute illness, activity-limiting unexplained illness or symptoms, or known active cancer) able to have a visit for 3–8 hrs of onsite data collection (depending on level of testing). All participants underwent a verbal review of the IRB-approved consent (Western IRB). Study results were returned to participants (within 10–12 weeks after the visit), who were encouraged to involve their primary care physicians. Additional details of this study can be found in Perkins et al., 2018^20^.

### Identification of Genetic Findings

Whole genome sequencing (WGS) was performed as described in Telenti et al., 2016^34^. Genetic variants were annotated using integrated public and proprietary annotation sources, including ClinVar^35^ and HGMD Professional^36^. To identify potentially medically significant rare monogenic variants, we used an internal version of HLI Search in a two-step process: the first step focused on allele frequency <1% in the HLI cohort with annotation of pathogenic/likely pathogenic or disease mutations using ClinVar^35^ and HGMD Professional^36^ as well as predicted loss of function variants. A second query of the patients’ whole genome sequencing data was targeted to reported disease causative genes/variants where phenotypes collected through our full range of tests suggested a genetic basis for disease. We developed disease-specific panels, using ClinVar^35^, HGMD^36^, HPO^37^, and OMIM^38^ databases, based on participants’ diagnoses. Common pathogenic variants with allele frequency >1%, including the *CFTR* p.Phe508del, *GJB2* p.Met34Thr, *CYP21A2* p.Val282Leu, *SERPINA1* p.Glu366Lys and p.Glu288Val, *F2* c.*97G>A, *FLG* p.Ser761fs and p.Arg501*, *F5* p.Arg534Gln, *G6PD* p.Val68Met, *BTD* p.Asp446His, and *HFE* p.Cys282Tyr were also reported.

Monogenic rare variants were manually interpreted using ACMG guidelines^39^ and were classified as pathogenic, likely pathogenic, variant of uncertain significance (VUS), or benign. The HLI database, which integrates allele frequencies for variants derived from >12,000 individuals with phenotypes, was also employed for variant interpretation. Two pairs of clinical geneticists and research scientists evaluated genomics variants and corresponding clinical data for each case. For expressivity analysis, clinical presentations listed in OMIM clinical synopses or in publications such as GeneReviews or primary studies were employed. Clinical geneticists, research scientists, and the clinical team discussed reportable variant of uncertain significance (VUS-R) classification based on participants’ genetic and clinical data.

### Identification of Short Tandem Repeat

Short tandem repeats (STRs) was identified using TREDPARSE software as described previously in Tang et al., 2017^40^.

### Metabolomics

The non-targeted metabolomics analysis of 1,007 metabolites was performed at Metabolon, Inc. (Durham, North Carolina, USA) on a platform consisting of four independent ultra-high performance liquid chromatography-tandem mass spectrometry (UPLC-MS/MS) methods. The detailed descriptions of the platform can be found in our previous publications^19,20^. The blood tests for prediabetes, Quantose impaired glucose tolerance (IGT) and insulin resistance (IR) were also performed^41,42^. Blood plasma was used for analysis, and values from multiple experimental batches were normalized into Z-scores based on a reference cohort of either 42 (n=456) or 300 (n=289) self-reported healthy individuals run with each batch.

The 42- and 300-normalized batches were converted to the same scale using linear transformation based on the values obtained from 7 runs that included both the 42 and 300 controls. Samples with metabolite measurements that were below the detection threshold were set as the minimum value for that metabolite. Other missing metabolite levels were imputed using the missForest R package.

To expand our analyses, we included metabolome data from 1,969 European ancestry twins enrolled in the TwinsUK registry, a British national register of adult twins^43^. We previously reported a detailed study of the genetic variants influencing the human metabolome in this cohort^19^. For the present study, we used data from serum samples that were collected at the first of three visits, when the ages of participants ranged from 33 to 74 years old (median 51). The cohort is mainly composed of females (96.7%), and the sample set included 388 monozygotic twin pairs, 519 dizygotic twin pairs, and 155 unrelated individuals. In common with the Health Nucleus cohort, the non-targeted metabolomics analysis of 901 metabolites was performed at Metabolon, Inc. (Durham, North Carolina, USA). As before, serum was used for analysis, and the resulting raw values were transformed to z scores using the means and standard deviations.

### Gene-based collapsing analysis

We performed a gene-based collapsing analysis to identify genes where carrying a rare functional variant was associated with increased or decreased metabolite levels relative to non-carriers. For this purpose, “rare functional variant” was defined as having a minor allele frequency (MAF)<0.5%, a CADD score>15, and being annotated as coding. These carrier statuses for each gene were used as the “genotypes” in genetic association analyses. A genetic similarity matrix (GSM) was constructed from ~300,000 variants that represented a random 20% of all common (MAF>5%) variants genome-wide after linkage-disequilibrium (LD) pruning (r2 less than 0.6, window size 200 kb) and was used to model the random effect in the linear mixed model via a “leave-out-one-chromosome” method for each tested variant. Each gene was tested independently using customized Python scripts wrapping the FaST-LMM package^44^. Analyses were performed both in all participants and separately in European ancestry participants only. The phenotypes analyzed were the levels of each metabolite, which were rank-ordered and forced to a normal distribution within each cohort before being combined for the genetic analysis. Statistically significant associations were identified using Bonferroni correction for multiple tests. Test statistic inflation was assessed with QQ plots.

### Cholesterol to HMG Ratio

The cholesterol/HMG ratio of each individual was calculated as the normalized percentile (using the pnorm command in R). Cholesterol metabolite value divided by the normalized percentile HMG metabolite value.

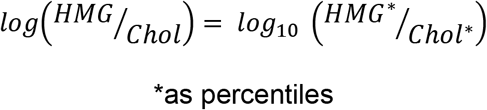

### Non-Contrast Whole-Body MRI

Study participants underwent non-contrast whole-body MRI (GE Discovery MR750 3.0T or Siemens Skyra 3.0T) using protocols and quantitative postprocessing for volumetric brain imaging (Neuroquant, CorTechs Labs prospectively and an in-house algorithm retrospectively), brain white-matter assessment, whole body detection of solid tumors with diffusion-weighted imaging^25, 45^, neurological and cardiovascular assessment, cardiac structure and function, liver-specific fat and iron estimation, and quantitative body compartment-specific fat and muscle estimation (Advanced MR AnalyticsAB) (Supplemental Table S1). The total time in the scanner was 60-90 minutes depending on the scanner and patient height. On the GE Discovery, coil configuration comprised a 16-channel head and neck coil, 8-channel cardiac coil, and 32-channel body array coil. On the Siemens Skyra, coil configuration comprised 64- or 20-channel head and neck coil (depending on the subject’s head size), 18-channel body array coil, 30-channel body array coil, and 32-channel spine coil. Also, for scanning on the Siemens Skyra we used the Abdomen DOT and Cardiac DOT engines. MRIs were interpreted by three radiologists (authors D.K. and N.H.; T.F., acknowledgements), a neuroradiologist and two other radiologists with fellowships in body imaging. All three radiologists had prior experience and training in interpreting imaging across the whole body including the brain. A quality control program was in place where a subset of studies was overread by a separate radiologist.

### Cardiac Tests

We used five different cardiac tests: cardiac computed tomography (CT), echocardiogram, 12-lead hospital grade electrocardiogram (ECG), cardiac rhythm monitoring, and cardiac CINE MRI. All cardiac test results were interpreted by a board-certified cardiologist (author A.M.K.) See Supplemental Table S2 for abnormal criteria. CT of the chest (heart and lungs, lung apices were not imaged) data were acquired for 671 individuals using a 64 slice GE Revolution CT (GE Healthcare, Milwaukee, WI). Gated axial scans with 2.5 mm slice thickness were performed using a tube energy of 120 kVp and the tube current adjusted for subjects’ body mass index. Images were subsequently analyzed using an A-W Workstation (GE Healthcare, Milwaukee, Wisconsin) and regions of coronary calcification were manually identified and Agatstan scores were computed. Data from two-week cardiac rhythm monitoring (Zio XT Patch; iRhythm Technologies, Inc., San Francisco, CA) kits were applied during the study visit or provided with instructions for home application were available for 915 individuals. ECG data were available for 876 individuals. Echocardiographic data were acquired using a Vivid* E95 Ultrasound System (GE Healthcare, Milwaukee, WI). Subjects underwent complete echocardiographic studies including M-mode, 2-D grey scale, pulse and color Doppler imaging.

### Dual-Energy X-Ray Absorptiometry

Dual-Energy X-ray Absorptiometry (DEXA) was conducted using Lunar iDXA with Pro Package (GE Healthcare) and was used for skeletal and metabolic health assessment.

## RESULTS

### Study Population

The cohort was composed of self-referred individuals with a median age of 54 years (range 20-89+ yrs, 33.8% female, 70.6% European) between September 2015 and April 2018. All participants had a primary physician. Our precision health platform includes whole-genome sequencing (WGS), non-contrast whole-body magnetic resonance imaging (MRI)^22,23^ including cardiac MRI and non-contrast MRI/MRA of the brain, global metabolomics including a blood test for prediabetes (Quantose IR and IGT)^42,41^, additional blood tests including complete blood count, kidney and liver function tests, vitamin and hormone levels, a lipid panel, cancer tumor marker screening, heavy metal screening, and blood sugar, echocardiography^46^, ECG^29^, non-contrast coronary artery calcium scoring CT^47^, and 2-wk cardiac rhythm monitoring in an effort to identify age-related chronic disease risks associated with premature death. Family and personal histories were also collected from participants.

### Medically Significant Genetic Findings

Among our cohort, 24.3% of participants had MSFs that may contribute to the risk of age-related chronic conditions. Medically significant findings determined by ACMG criteria are summarized in Figure 1. A total of 206 unique medically significant variants in 111 genes were identified. Forty individuals (3.4%) had more than one MSF, and one individual had MSFs in four genes. The major categories of diseases include hematological (30%). cancer (28%), cardiovascular (20%), endocrine (5%) and neurologic disorders (2%). Details of each case and the corresponding clinical associations detected in our phenotypic tests are provided in the addendum (Supplemental Table S3). Frequently observed low to moderate penetrance variants, including *F2, F5*, and *ALDH2*, were found in 7% of this cohort. The most commonly affected genes (number of cases) were *CHEK2* (n=17), *MYBPC3* (n=8), *BRCA2* (n=7), *ATM* (n=6), *HOXB13* (n=6), and *LDLR* (n=6) for the autosomal dominant conditions. For the autosomal recessive conditions, the most commonly affected genes for MSFs (homozygous or potential compound heterozygous) were *HFE* (n=15), *BTD* (n=5), and *GJB2* (n=2). For the X-linked recessive conditions, we identified six cases with glucose-6-phosphate dehydrogenase (G6PD) deficiency. Thirty-two individuals (2.7%) had pathogenic or likely pathogenic variants identified in the ACMG 59 genes associated with potential high medical importance.

**Figure 1.**
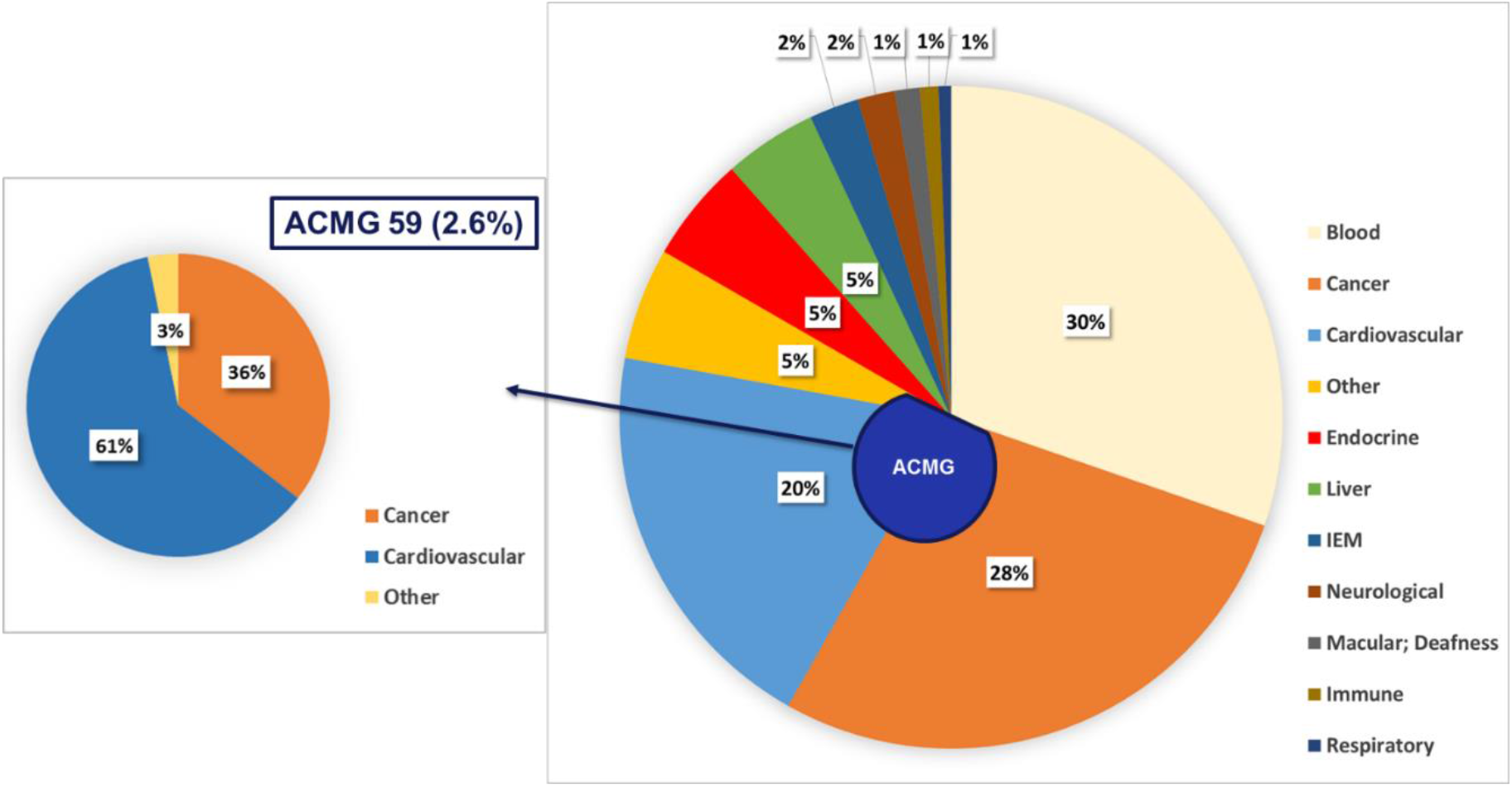
Medically Significant Genetic Findings by Disease Group. 24.3% of participants have medically significant genetic findings, including heterozygous for autosomal dominant conditions, biallelic or homozygous for autosomal recessive conditions, and hemizygous for X-linked conditions. Variants were manually interpreted using ACMG guideline^39^. Reportable variants of unknown significance comprised 4.8% of participants. Thirty-two individuals (2.7%) had pathogenic or likely pathogenic variants identified in the ACMG59 genes, a minimum list of genes to be reported as incidental or secondary findings^48^. IEM, inborn error of metabolism.

In addition to pathogenic and likely pathogenic variants as recommended in the ACMG guidelines^39^, a category of reportable variants of uncertain significance (VUS-R) was created. Given each participant had a full range of screening tests, a VUS finding may explain the disease diagnosis unique to a family with compelling clinical presentations^49^. As an illustration of the value of VUS-R, we identified a variant (c.237-1G>A) in the *ERLIN2* gene as a VUS for spastic paraplegia (MIM 611225), a neurological autosomal recessive disorder in a 65-year-old male with progressive gait difficulties and ataxia; his son (30s) had similar neurological symptoms. The initial clinical diagnosis was spinocerebellar ataxia (SCA). The WGS results did not detect single nucleotide variations (SNVs) or short tandem repeats (STRs) in genes involved with SCA disorders^50,47^. The MRI result did not reveal atrophy of the brainstem or cerebellum. A report by Rydning et al. showed a novel *ERLIN2* heterozygous missense variant in two unrelated families with an autosomal dominant inheritance of hereditary spastic paraplegias^51^. The c.237-1G>A variant was also identified in a 57-year-old male patient with suspected hereditary spastic paraplegia without affected family members (Praxis fuer Humangenetik Tuebingen, SCV000493455.5). The collective evidence led to an update of this variant as VUS-R for the autosomal dominant form of hereditary spastic paraplegias to this participant and his treating clinician to assist on the diagnosis of his neurological disorder.

To investigate the expressivity of MSF in our cohort, we examined the clinical phenotype associated with the MSF. Clinical synopses in OMIM and phenotypes listed in the published literature were employed for the analysis. Clinical associations, including family histories, personal histories, and phenotypic test results with genomic findings, were identified among 63.1% study participants (n=130) in individuals with high to moderate penetrance variants (n=206). Details of clinical associations with cancer, cardiovascular, cirrhosis, and endocrine conditions are listed in Table 1. Cardiovascular and endocrine diseases achieved substantial associations between MSFs and clinical phenotypes (89% and 72% respectively), indicating that our integrated approach is able to provide valuable information, enabling clinicians to integrate genetic analysis into their diagnostic assessments. Clinical associations were observed among 69.0% of study participants (n=40) with high and moderate penetrant inherited cancer variants, and family history contributed to a large fraction of the correlation. The integration of genetic variants with predictive value and comprehensive phenotypic tests can further refine the differential diagnosis, direct the use of therapies or surveillance screenings, and guide appropriate genetic counseling and testing of at-risk family members.

**Table 1.** Clinical Associations with Rare Monogenic Variant by Disease Group and Test. Clinical associations include past medical history, family history and results from clinical tests. Clinical synopses in OMIM and clinical phenotypes listed in the literature are used for clinical correlation analyses. The clinical features observed in individuals with ANGPTL4, APOC3 or NPC1L1 variants were protective effects (i.e., lowering blood lipids). *AR STR is a risk allele based on case-control study. Abbreviations: ECHO, echocardiography; MRI, magnetic resonance imaging; CT, computed tomography coronary artery calcium scoring; ECG, electrocardiogram. (n)=number of individuals

| Conditions | Genes | Total | Test Associations |
| --- | --- | --- | --- |
| **Cardiomyopathy, Arrhythmia, and others** | ANK2, CFHR5, DSC2 (x2), ENG, GPD1L (x2), KCNE2, KCNQ1 (x2), LMNA, MYBPC3 (x8), MYH7 (x5), MYL2 (x4), MYLK (x2), PKP2, RBM20, RYR2, SCN1B, SMAD6, TCAP, TNNT2, TTN (x2), TTR (x2), VCL | 42<br>(3.5%) | Family history (23);<br>ECHO (17); ECG (11) |
| **Dyslipidemia** | ANGPTL4, APOB (x5), APOC3 (x2), LDLR (x6), LPL (x3), MEF2A (x2), NPC1L1, PAFAH1B2, PCSK9 (x3) | 24<br>(2.0%) | Clinical tests (18);<br>Family history (12);<br>CT (10) |
| **Cirrhosis** | HFE (3 homozygous and 12 compound heterozygous) | 15<br>(1.3%) | MRI (7); Clinical tests (4); ECHO (1) |
| **Cancer** | AR STR (x2)*, APC, ATM (x6), BARD1, BRCA1, BRCA2 (x7), BRIP1 (x3), CHEK2 (x16), CYLD, EPCAM, FAM175A, FH (x2), GEN1, (x2), GLMN, HOXB13 (x6), LZTR1 (x4), MSH6, NBN (x4), NF1, PALB2, PMS1, PMS2, RAD50 (x2), RAD51C, RB1, RECQL, TP53 | 69<br>(5.8%) | Family history (38);<br>MRI (2) |
| **Endocrine** | ABCC8 (x2), FAAH, GLMN, HNF1A, HNF1B (x2), INSR, MC4R, NEUROD1, NOBOX, NR3C1, PCSK1 (x2), PPP1R3A (x2), PROK2, TBC1D4 | 18<br>(1.5%) | Family history (4),<br>MRI (4), Clinical tests (4) |

Each individualized autosomal dominant MSF enables the study of family member risk by their specific variants, thus, expanding the clinical utility of disease diagnosis. Figures 2A and 2B illustrate families with positive family histories of hyperlipidemia and breast cancer, respectively. Pathogenic/likely pathogenic variants were identified in the index case, and the clinical molecular genetic diagnosis was established. At-risk individuals from these two families were labeled in pink (n=10). This information can direct the clinician to provide genetic counseling and testing of other relatives that may be high-risk for heritable disorders.

**Figure 2A.**
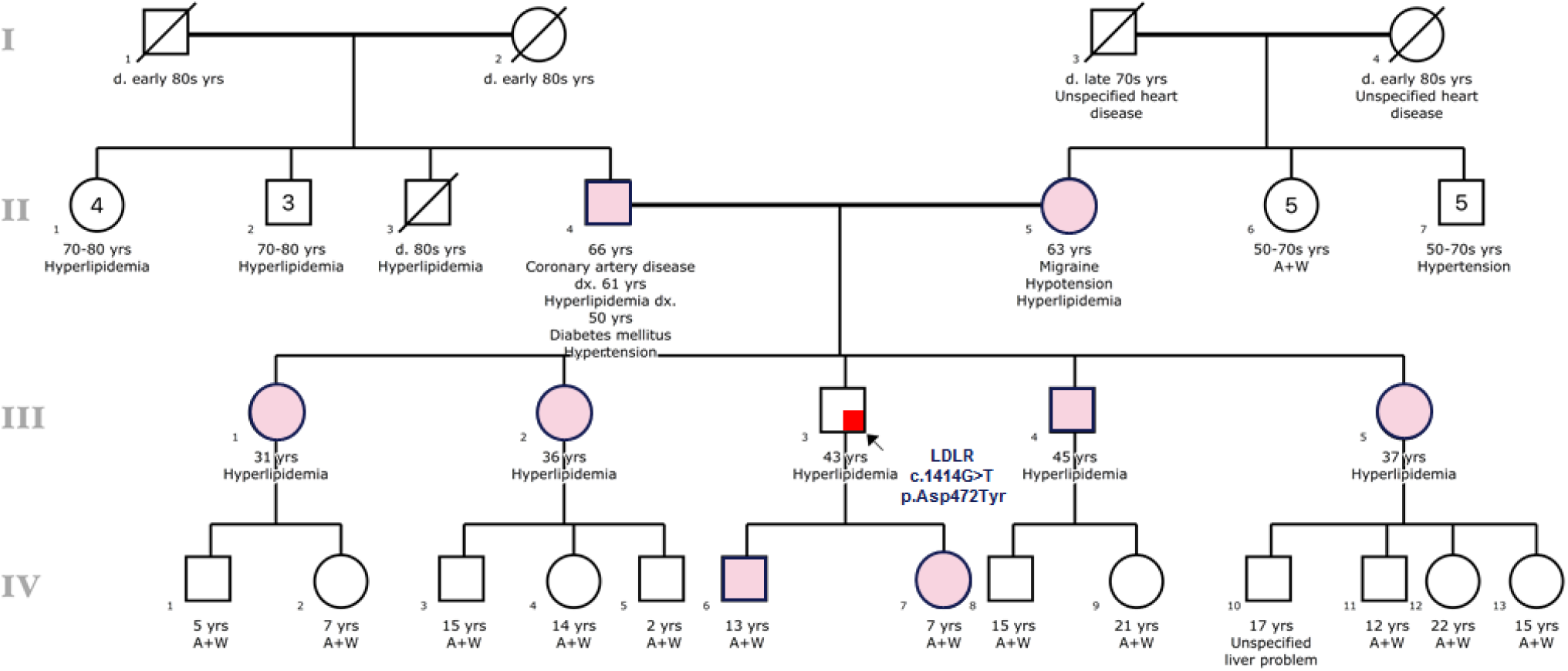
Pedigree of a Familial Hypercholesterolemia Individual. The LDLR c.1414G>T (p.Asp472Thr) variant, classified as likely pathogenic, was identified in the proband whose phenotypes were compatible with the clinical presentation of familial hypercholesterolemia. Family members highlighted in pink may benefit from cascade testing.

**Figure 2B.**
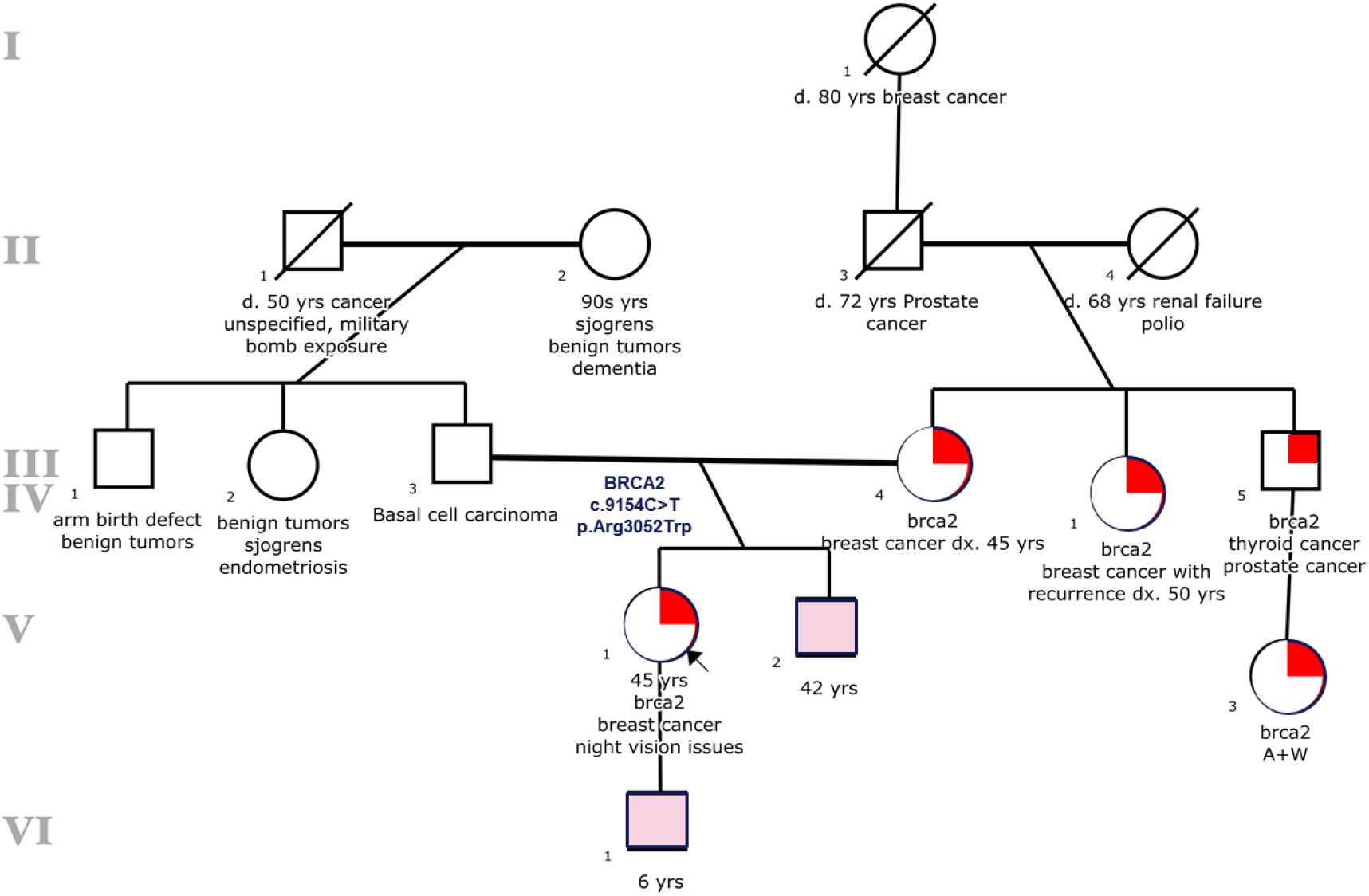
Pedigree of a Hereditary Breast and Ovarian Cancer Syndrome Individual. The BRCA2 c.9154C>T (p.Asp3052Trp) variant, classified as pathogenic, was identified in the proband whose family history and personal history were compatible with the clinical presentation of hereditary breast and ovarian cancer syndrome. Family members highlighted in pink may benefit from cascade testing.

We have selected three cases to illustrate the utility of genome sequencing to assess a wide range of genetic variants for diagnostic purposes: Case 1. A diagnosis of cystic fibrosis (CF) was made on a male participant (mid-50s). Neither this participant nor his physician had a clinical suspicion of CF and this participant has a history of digestive symptoms and chronic sinus infections. The imaging findings found tree-in-bud nodularity at the right mid and upper lung zone and mild bronchial wall thickening (Figure 6G). Two variants, c.3846G>A (p.Trp1282*) and c.3454G>C (p.Asp1152His), were identified in the *CFTR* gene. Results from genetics and clinical presentation were consistent with atypical cystic fibrosis. The detection of p.Asp1152His (D1152H) variant prompted this client to be referred to a local cystic fibrosis center for consideration of treatment with Ivacaftor and/or pancreatic enzyme supplements^52,53^.

Case 2. An adult (60s) was found to have a pathogenic variant (c.264C>A, p.Asn88Lys, homozygous) in the *RAPSN* gene suggesting a diagnosis of the congenital myasthenic syndrome. He presented with mild extraocular muscle weakness. His brother (60s) has been treated for myasthenia gravis for over two decades. The clinical presentation of the congenital myasthenic syndrome is similar to that of myasthenia gravis^54^. The same genetic variant was identified in his brother, thus correcting his diagnosis to the congenital myasthenic syndrome. He has now enrolled in a phase III clinical trial, and treatments of immunosuppressive drugs and intravenous immunoglobulin therapy have been terminated.

Case 3. A participant (70s) arrived at our clinic asymptomatic for a routine results review and suddenly felt lightheaded and dizzy and was found to be bradycardic with heart rate of 30 beats per minute. Persistent symptoms led staff to call 911 and the participant was hospitalized. A cardiovascular evaluation revealed symptomatic 2nd-degree heart block and a dual chamber pacemaker was implanted. WGS identified c.398G>A (p.Arg133Gln), a likely pathogenic variant in the *LMNA* gene, associated with the LMNA-related disorder. This participant had evidence of left ventricular hypertrophy as assessed by ECHO, arrhythmia, and symptoms of fatigue, shortness of breath, and fainting episodes (syncope), as well as a family history of sudden death in three 2nd degree relatives (70s to 80s), consistent with the molecular genetics diagnosis of LMNA-related dilated cardiomyopathy (MIM 115200)^51^. The variant was also detected in his daughter (40s), currently without clinical manifestations. Both participants were advised to follow-up with a cardiovascular genetics specialist for actionable clinical intervention recommendations.

These three cases illustrate previously undetected genetic diagnoses with a potential therapy initiation, correction of misdiagnosis and incorrect therapy, and an actionable implementation of a medical procedure for the intervention.

### Autosomal Recessive Genetic Findings

In our cohort, 86.3% of individuals had at least one autosomal recessive (AR) variant in 680 genes. The most commonly detected carriers (number of cases) were *SERPINA1* (n=99), *FLG* (n=95), *BTD* (n=84), *HFE* (n=84), and *GJB2* (n=71) (Table 2). The maximum number of AR variants per individual was eight (n=3, unrelated). Details of AR variant findings (>1% observed frequency) are provided in Table 2 including eighteen genes not commonly found in gene panels for carrier screens.

**Table 2:**
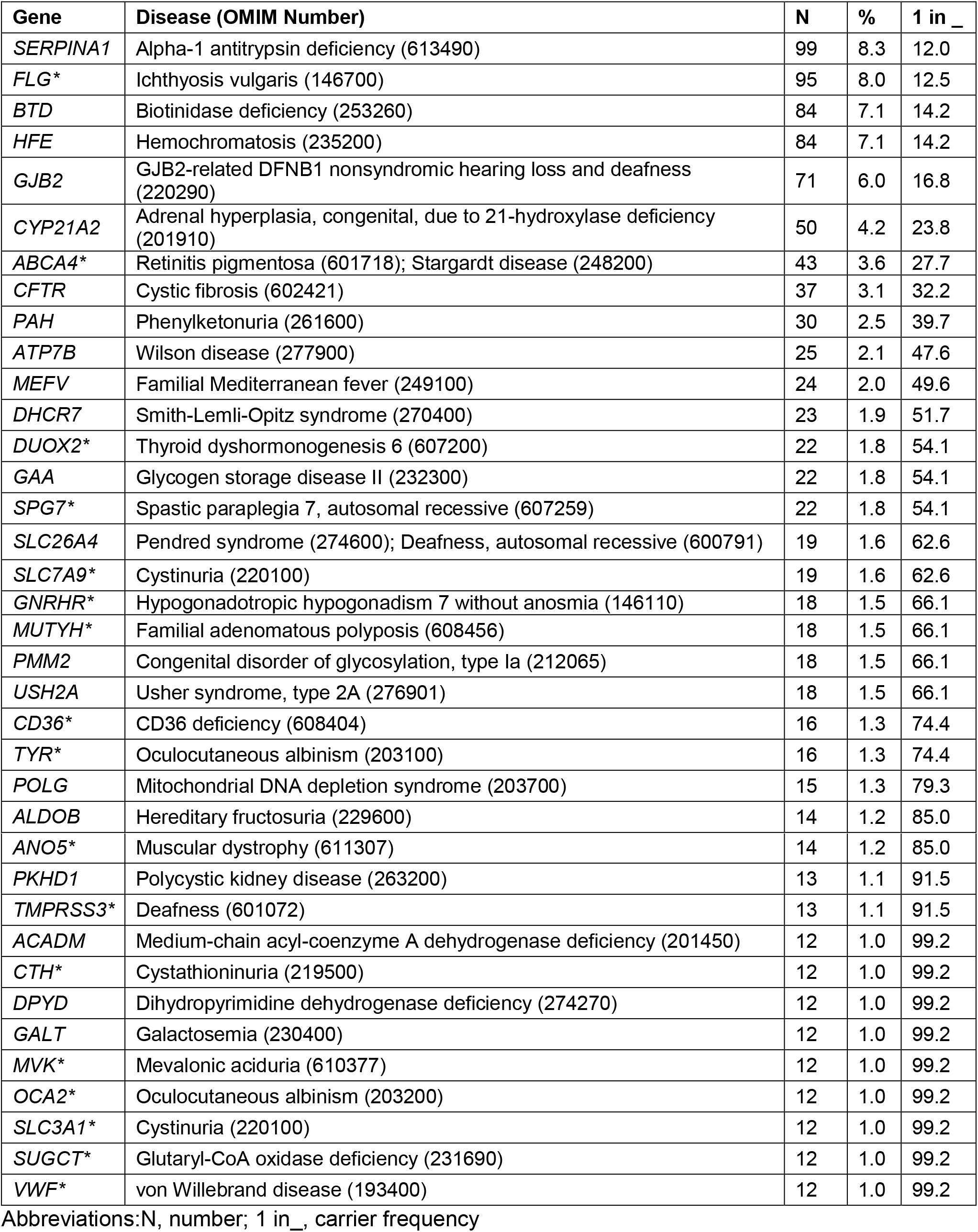
Frequency of Autosomal Recessive Carriers (*not commonly included in carrier screens)

From our study, we observed that carriers of known AR conditions had penetrance for imaging abnormalities and metabolomic biomarkers consistent with the expression of the AR phenotype. For example, pathogenic or likely pathogenic variants in the *PKHD1* gene, the causative gene for autosomal recessive polycystic kidney disease (ARPKD, MIM 263200) were identified in thirteen individuals. Eight of the carriers (average age 53.1 yrs) had numerous liver and kidney cysts detected by MRI consistent with a previous report that adult *PKHD1* carriers present clinically-isolated polycystic liver disease^55^. The remaining five *PKHD1* carriers (average age of 35.6) had no detectable liver or kidney cysts. Eighteen percent (12/68) of heterozygotes for the *HFE* p.Cys282Tyr variant have high R2*, a marker of liver iron content, compared to only 8% (87/1034) in individuals with normal genotype (p=0.0156, Fisher Exact), suggesting iron regulation is compromised^56^. For metabolic penetrance, ten out of thirty (33%) of *PAH* carriers had elevated phenylalanine detected by metabolomics^57^. Three out of four *ALPL* carriers had reduced serum alkaline phosphatase. One *ALPL* carrier (p.Phe328del) with reduced serum alkaline phosphatase was also found to be osteopenic in the left and right femur and lumbar spine detected by dual-energy x-ray absorptiometry (DEXA)^58^.

The associations between gene variants and metabolic penetrance are shown in Figure 3 and Supplemental Tables S4 and S5. These results were obtained by the integration of HLI search and biochemical pathways. We searched for an intersection of sequence variants predicted to be loss of function (LOF) and metabolites found to be at least six standard deviations (±6S.D.) from the normal range. A fraction of the detected metabolites that were extremely elevated was derived from medications, over-the-counter medications, and supplements metabolites. These findings are not discussed in this report. Two approaches were employed to investigate the associations between genomic variation and metabolite level. The biochemical pathways of individuals with extremely elevated metabolites were investigated for candidate genes. Then, an in-depth analysis was performed to identify rare functional variants that might be causal. For the other approach, we evaluated the metabolite penetrance in our cohort. For individuals who carried pathogenic or likely pathogenic variants associated with metabolic conditions, we analyzed the metabolic penetrance using metabolic features and laboratories abnormalities listed in the OMIM synopsis. Both approaches were able to establish associations (15% and 13%, respectively) and details are listed in Figure 3 and Supplemental Table S4B. An interesting case illustrating an association was a LOF nonsense variant, c.13C>T (p.Arg5*), in the *TTPA* gene, associated with ataxia with isolated vitamin E deficiency (MIM 277460). Three family members had the maternally segregated variant, and all had markedly reduced levels of vitamin E, ranging from -3.6 to -6 S.D., suggesting a high degree of penetrance. Clinical ataxia was not observed in these participants.

**Figure 3.**
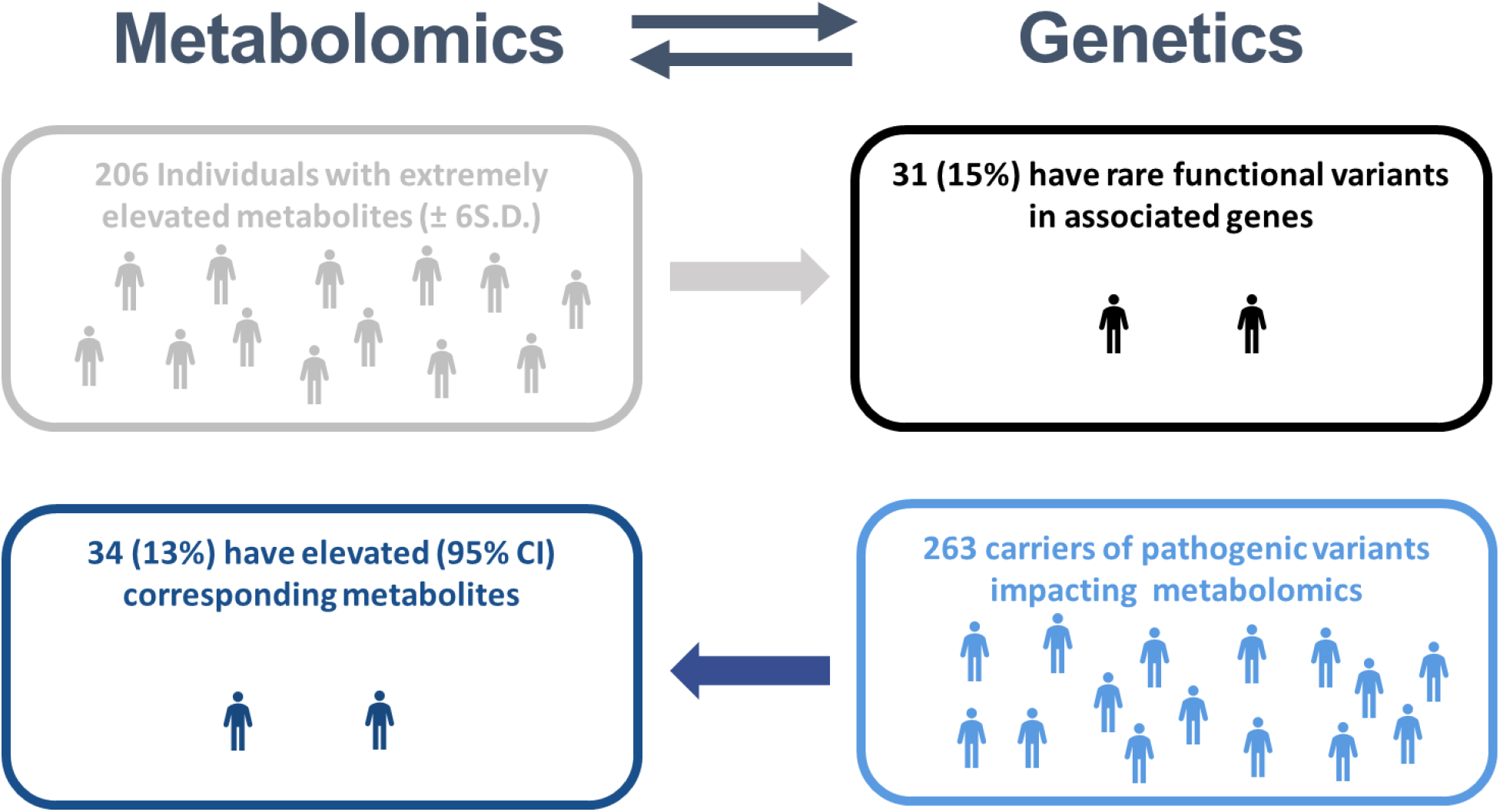
Association of Genetic Variants and Metabolites. Thirty-one out of two hundred and six individuals (15%) with extremely elevated or decreased metabolites (±6S.D.) had rare functional variants in pathway-associated genes (Figure 3, Supplemental Table S4A). Thirty-four carriers (13%) of pathogenic or likely pathogenic variants identified by WGS that were found in genes associated with metabolic pathways had either elevated or decreased (95% CI) levels in the corresponding metabolites detected by metabolomics.

To identify novel associations between genomic variants and metabolites, we used a gene-based collapsing analysis to identify genes where rare functional variants were associated with a statistically significant difference in the levels of any of the 1,245 measured metabolites. We began by focusing on 153 genes that are used in newborn screening and that are known to cause inborn errors of metabolism. We identified significant associations between the *PAH* gene (phenylketonuria) and the metabolites phenylalanine and gamma-glutamylphenylalanine as well as between the *ETFDH* gene (glutaric academia) and octanoylcarnitine, decanoylcarnitine, and the nonanoylcarnitine. Expansion of analysis genome-wide identified 19 significant associations with 11 other genes (Supplemental Table S5). The associations identified known conditions, such as dimethylglycine dehydrogenase deficiency and histidinuria. However, five novel associations were identified: 1.) between the carbohydrate 1,5-anhydroglucitol (1,5-AG) and the *SLC5A10* gene; 2.) between the amino acids alpha-hydroxyisovalerate and 2-hydroxy-3-methylvalerate and the *HAO2* gene; 3.) between the amino acid 5-hydroxylysine and the *HYKK* gene; 4.) between N-acetyl-beta-alanine and the *PTER* gene; and 5.) between alpha-ketoglutaramate and the *NIT2* gene. *NIT2* is known to break down alpha-ketoglutaramate, and we have identified loss of function variants.

### Short Tandem Repeat Findings

Short Tandem Repeat (STR) measurements utilized our previously reported algorithm for the 40 disease genes caused by repeat expansions^47^. We identified a spinocerebellar ataxia case (SCA17) with repeat expansion causative of the participant’s gait abnormality. Details of participants with intermediate-length or full-penetrance STR expansions are provided in Supplemental Table S6. The number of STRs in the androgen receptor (AR) gene has been inversely correlated with the gene’s transcriptional activity^59^. The association of the *AR* CAG repeats with prostate cancer and benign prostatic hyperplasia have been studied extensively, and results have been elusive and controversial^60,61,62^. The CAG repeat sequence ranges normally from about 8 to 31 repeats with an average of about 20 STR in the *AR* gene^63^. The analysis of our male cohort of 5234 samples showed that STR lengths in the *AR* gene had different distributions and mean lengths (21.9 v.s.19.8) between European Americans and African Americans respectively. The group of individuals with STR ≤17 was 25.5% in African Americans compared to 4.3% in European American (p<0.00001). In addition to 1190 subjects present here in this report, we further analyzed the STR length in the *AR* gene for all male HN clinic participants (n=1002). In the case-control study with a repeat length of ≤17 versus >17, we found an approximately 4-fold increased risk of prostate cancer (OR 3.72, 95% CI [1.37-10.07]; t-test p=0.01). We found an approximately 2-fold increased risk of prostate disease including enlarged prostate and prostate cancer (OR 1.92, 95% CI [1.02-3.64]; t-test p=0.04). Individuals with STR ≤17 had increased prostate size measured by MRI (mean 32.21 cm3, p=0.046) compared to individuals with longer STRs (mean 27.85 cm^3^) (Figure 4).

**Figure 4:**
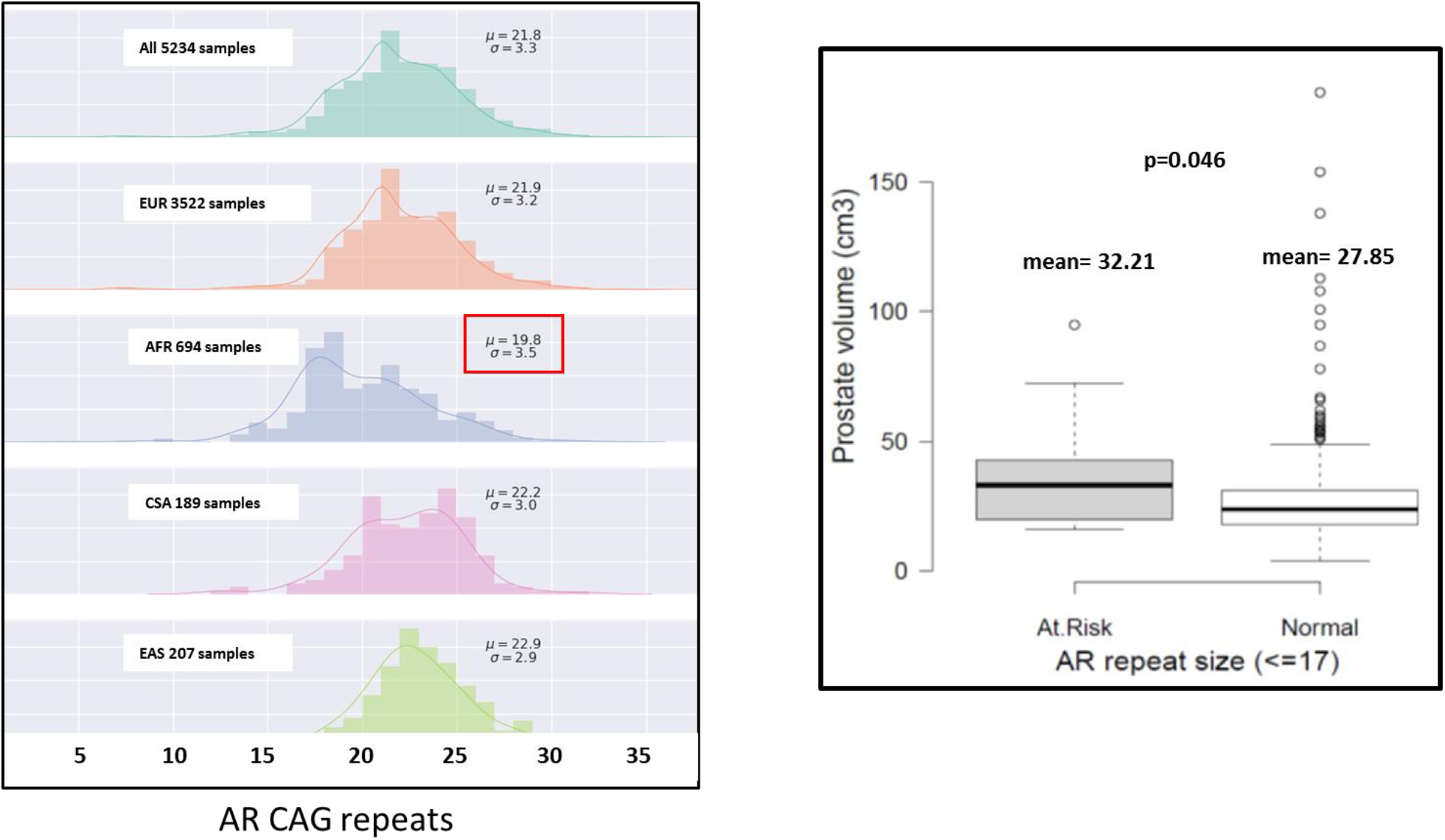
Association of the *AR* CAG Repeats with Prostate Volume and Population Frequency of the CAG Repeats in the Androgen Receptor (AR) Gene. The analysis of our male cohort of 5234 samples showed that STR lengths in the *AR* gene had different distributions and mean were displayed. Individuals with STR (<=17) had increased prostate volume measure by WB-MRI (mean=32.21cm^3^) compared to individuals with longer STRs (mean=27.85cm^3^). Individuals taking finasteride, dutasteride, or tamsulosin were not included in this analysis. EUR, European American; AFR, African American; CSA, Central South Asian; EAS, East Asian.

### A Metabolomics Approach for Personalized Drug Response

A functional approach using metabolomics was employed to measure the efficacy of treatments for gout and hypercholesterolemia/dyslipidemia. Inhibitors of xanthine oxidase, such as allopurinol, are commonly used in the treatment of diseases are associated with high levels of uric acid (i.e., urate), including gout and tumor lysis syndrome. Metabolite levels of xanthine, hypoxanthine, orotidine, orotic acid (i.e., orotate), and urate were employed for the analysis of four groups of individuals (Figure 5A). Individuals taking allopurinol (n=12, shown in blue) had elevated xanthine and orotidine levels compared to control individuals with carriers with *XDH* variants, associated with xanthinuria (MIM 278300). The individuals taking higher doses of allopurinol (200-300 mg, in red circle) had elevated xanthine and orotidine levels compared to individuals on the lower dose (50-100mg, in the black circle), indicating a dose-dependent effect (Figure 5A). The use of metabolomics enables physicians to monitor drug efficacy more precisely. Although urate as an indication of drug efficacy has been used in clinics for many years, the inclusion of precursors such as xanthine and hypoxanthine or off-target indicators such as orotic acid and orotidine may have the potential to aid and guide the safety and effective use for this important class of drugs. Low doses of allopurinol were associated with lower levels of xanthine and orotidine, indicating that the efficacy of the drug may not be maximized.

**Figure 5A.**
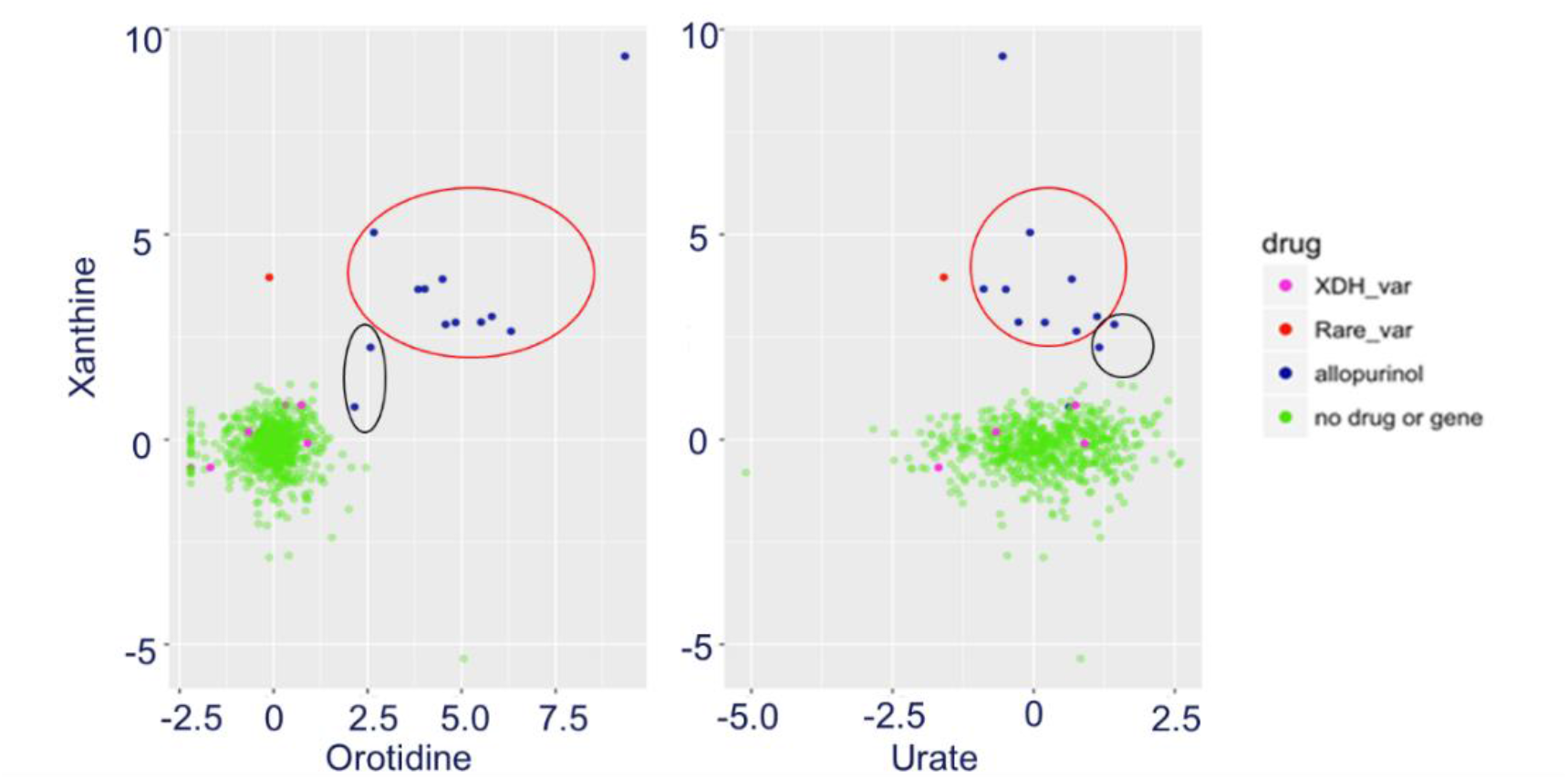
Measurement of Allopurinol Therapy Effect. Individuals taking allopurinol (n=12, shown in blue) had elevated xanthine and orotidine levels compared to control individuals. Individuals taking higher doses of allopurinol (200-300 mg, circled in red) had elevated xanthine and orotidine levels compared to individuals on lower dose (50-100mg, circled in black).

XDH_var: *XDH* variants, associated with xanthinuria; Rare_var: rare *XDH* variants in a suspected xanthinuria individual.

Cholesterol secretion, lipoprotein receptor activity, and *de novo* cholesterol biosynthesis are important for cellular cholesterol homeostasis^64^. Inhibition of hydroxy-3-methylglutaryl coenzyme-A (HMG-CoA) reductase has been employed to reduce low-density lipoproteins (LDL), one of the major risk factors for coronary artery disease (CHD)^65^. To measure the efficacy of HMG-CoA inhibitors such as statins, we measured the hydroxy-3-methyl glutarate (HMG) levels and calculated the ratio of cholesterol to HMG (CHO: HMG) for each individual. Individuals on statin therapy (n=52) had a lower CHO: HMG ratio (mean=0.85, p=0.001) compared to 603 individuals without any cholesterol lowing therapy (mean=1.17). Similarly, individuals on ezetimibe therapy (n=9) had a lower CHO: HMG ratio (mean=0.80, p=0.02) compared to individuals without any therapies, possibly through the upregulation of *LDLR* gene and protein expression^66^,^67^. Two individuals treated with the *PCSK9* monoclonal inhibitor exhibited lower ratios (0.4 and 4.6 percentile respectively). Two of our participants with pathogenic/likely pathogenic variants in the low-density lipoprotein receptor *(LDLR)* gene, indicating defects in the expression of *LDLR* or disruptions of the receptor’s ability to remove LDL from the blood, exhibited higher ratios (94^th^ and 96^th^ percentile respectively, Figure 5B). We suggest that the measurement of HMG and the ratio of CHO: HMG may be useful to clinicians and pharmaceutical companies to optimize the cholesterol-lowering therapy for high-risk individuals. Individuals with high and low ratios also are being studied to determine the possible genetic basis for our speculated increased rate of *de novo* cholesterol biosynthesis and possible effects on reducing the risk of atherosclerotic cardiovascular disease.

**Figure 5B.**
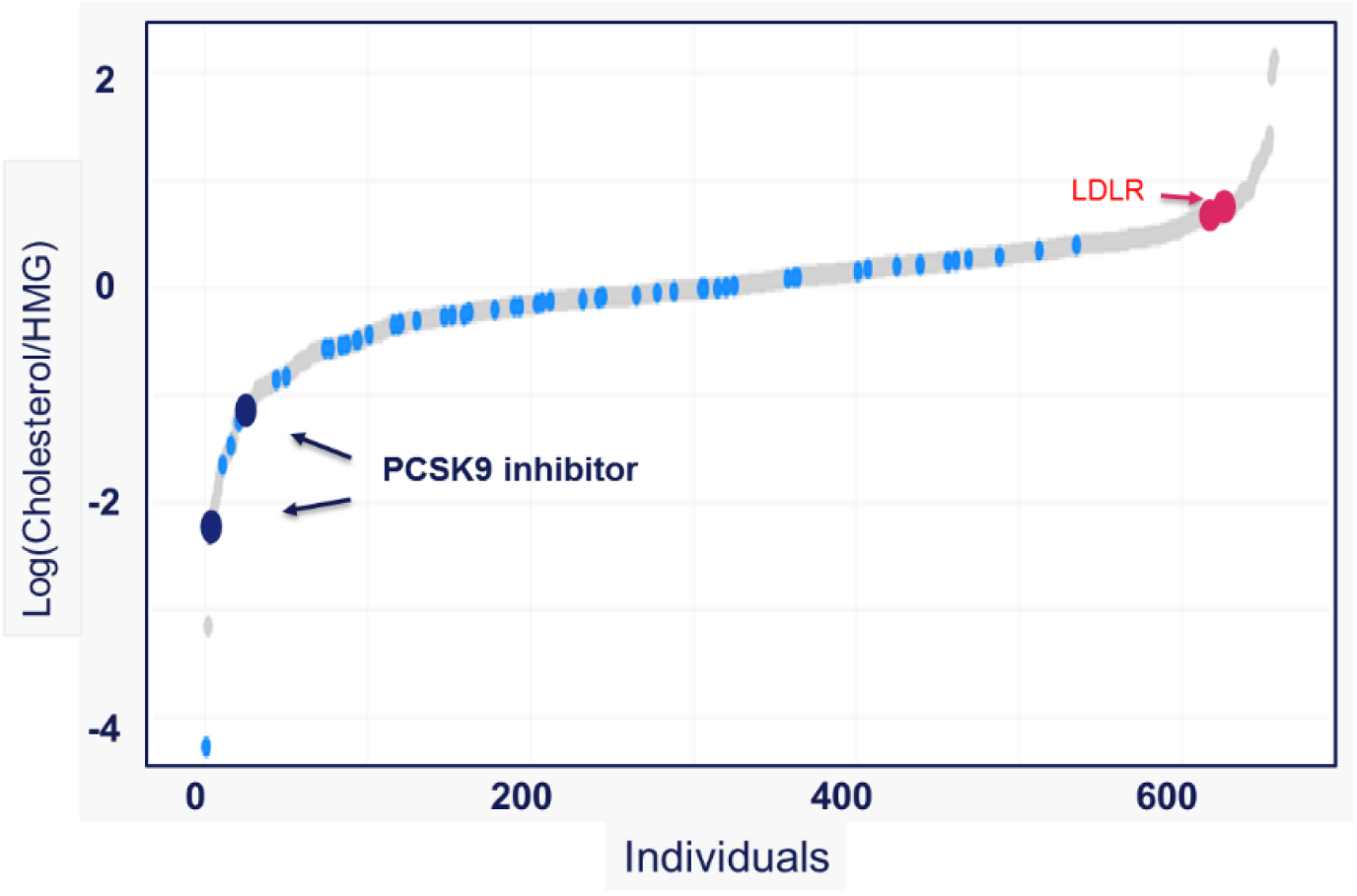
Measurement of Statin Therapy Effect Using Hydroxy-3-methylglutarate (HMG). Individuals on statin therapy (light blue dots) had a lower CHO:HMG ratio compared to individuals without any therapy (p<0.001). Individuals treated with a *PCKS9* inhibitor exhibited lower ratios (dark blue dots, p=0.002) and *LDLR* carriers exhibited higher ratios (red dots, p<0.03).

### Whole-body MRI Findings

While whole-body magnetic resonance imaging (WB-MRI) has been available for some time, only recently has it become clinically practical. Reduced acquisition times and a reduction in acquisition costs have facilitated the clinical introduction of this technique as a practical diagnostic tool.

WB-MRI is gaining popularity in detecting a wide range of diseases, including lymphoma, multiple myeloma, and systemic diseases such as diabetes, atherosclerosis and systemic musculoskeletal diseases^27^. The protocols used in this study comprise both conventional and research sequences, resulting in a comprehensive overview of a participant’s body and health status, including quantitative and qualitative analysis of the brain, neurological, cardiovascular, and liver disease, body composition, and tumor detection (Supplemental Table S1). Importantly, the primary purpose of the whole-body imaging in this study was two-fold: 1.) to derive quantitative imaging biomarkers which can be integrated with genetic and blood biomarkers for disease risk prediction and; 2.) to assess the participant’s current health status.

A diversity of disease is detectable via WB-MRI. Cerebrovascular disease, such as stenosis or aneurysms, detected through the use of 3D TOF non-contrast MR angiography to generate images of arteries in the brain. Regional brain atrophy, particularly in the temporal lobes, quantified using segmented 3D structural T1-weighted images of the brain. Quantitative biomarkers of an individual’s metabolic health can be obtained via segmentation of the body in to compartments such as lean muscle, visceral adipose tissue, and subcutaneous adipose tissue utilizing whole body Dixon imaging. Dixon imaging can discriminate hydrogen protons from water within muscle tissue and lipids in adipose tissue with high accuracy. Detection of tumor tissues is enhanced by the use of whole body diffusion-weighted imaging (DWI) which enables probing the underlying structure of tissue at a cellular level via the movement of water molecules distinguishing tissues with high cellularity from those that do not. Cardiac structure and function can be evaluated utilizing short and long axis views of the heart obtained using Cardiac CINE^45, 68^.

Overall, within this study, WB-MRI yielded findings resulting in a recommendation for repeat evaluation in 22% of the individuals in this cohort. Significant findings included 9.3% with elevated R2* (indicative of high levels of iron in the liver), 29.2% with elevated levels of fat in the liver indicative of fatty liver, 1.2% with a brain aneurysm, 1.3% with a body aneurysm (the majority were ascending aortic aneurysms), and 2.0% with a low hippocampal occupancy score relative to population (which can be due to medial temporal lobe atrophy that may be seen in neurodegenerative disease such as Alzheimer’s dementia). Furthermore, there were 2% of individuals with newly identified tumors. Except for brain meningioma, all cancers were confirmed by biopsy, post-resection tissue, or contrast-enhanced imaging prior to ablation. Within this cohort, we are not aware of any false positive (FP) biopsy findings. We considered findings for which follow-up biopsies were negative to be confirmed FPs. Including this cohort and all other patients who have visited our clinic to date (2850 in total), we know of only two patients (prostate and mediastinal lymph node) for whom follow-up biopsies were benign (Table 3). Furthermore, the average stage of tumors identified through our WB-MRI was early with fair prognosis, thus, increasing the chances of survival.

**Table 3:**
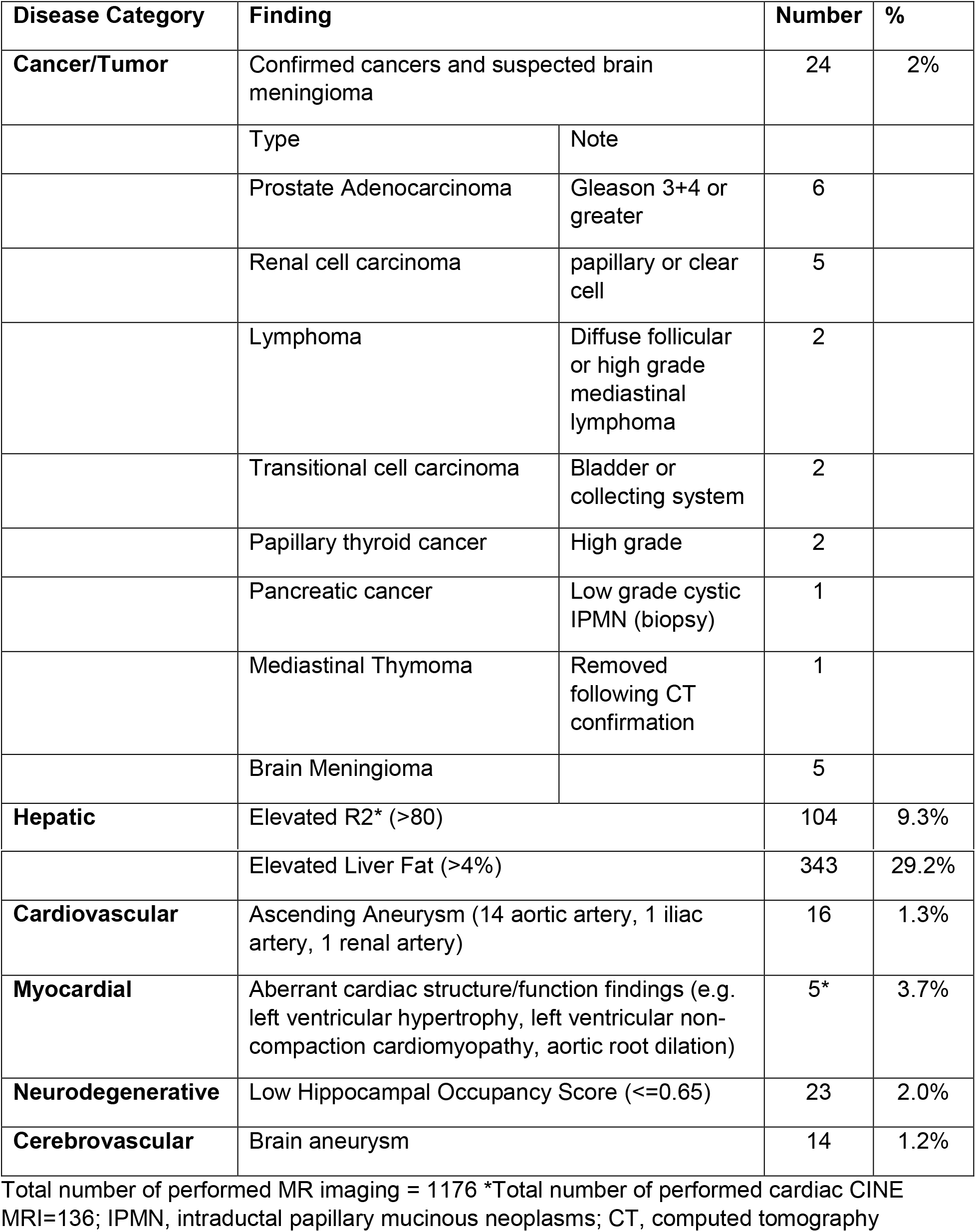
Phenotypic MR Imaging Findings by Disease Category.

| Disease Category | Finding |  | Number | % |
| --- | --- | --- | --- | --- |
| **Cancer/Tumor** | Confirmed cancers and suspected brain meningioma |  | 24 | 2% |
|  | Type | Note |  |  |
|  | Prostate Adenocarcinoma | Gleason 3+4 or greater | 6 |  |
|  | Renal cell carcinoma | papillary or clear cell | 5 |  |
|  | Lymphoma | Diffuse follicular or high grade mediastinal lymphoma | 2 |  |
|  | Transitional cell carcinoma | Bladder or collecting system | 2 |  |
|  | Papillary thyroid cancer | High grade | 2 |  |
|  | Pancreatic cancer | Low grade cystic IPMN (biopsy) | 1 |  |
|  | Mediastinal Thymoma | Removed following CT confirmation | 1 |  |
|  | Brain Meningioma |  | 5 |  |
| **Hepatic** | Elevated R2* (>80) |  | 104 | 9.3% |

|  |  |  |  |
| --- | --- | --- | --- |
|  | Elevated Liver Fat (>4%) | 343 | 29.2% |
| **Cardiovascular** | Ascending Aneurysm (14 aortic artery, 1 iliac artery, 1 renal artery) | 16 | 1.3% |
| **Myocardial** | Aberrant cardiac structure/function findings (e.g. left ventricular hypertrophy, left ventricular non-compaction cardiomyopathy, aortic root dilation) | 5* | 3.7% |
| **Neurodegenerative** | Low Hippocampal Occupancy Score ($\leq 0.65$) | 23 | 2.0% |
| **Cerebrovascular** | Brain aneurysm | 14 | 1.2% |

Total number of performed MR imaging = 1176 \*Total number of performed cardiac CINE MRI=136; IPMN, intraductal papillary mucinous neoplasms; CT, computed tomography

The integration of advanced imaging, clinical tests, and genomic sequencing achieved the clinical diagnosis of diseases which were not established in prior visits (Figure 6). For example, the detection of a pathogenic variant in the *NF1* gene (*de novo*) in an individual with optic glioma, white matter lesions, and stenosis in the middle cerebral artery (Figure 6A and 6B) allowed the clinician to confidently assess the detected variant as the cause of disease in this individual. Results from WB-MRI provided comprehensive clinical presentations of neurofibromatosis for this individual. Similarly, for individuals with the *PKD1, HNF1B, CFTR, MYH7*, and *KCNQ1* variants (Figure 6E, 6F, 6G, 6J, and 6M), WB-MRI, CT and ECG established the clinical-molecular diagnosis of inherited diseases. For the individual with the *HFE* p.Cys282Tyr and p.His63Asp variants, WB-MRI allowed the surveillance of disease progression on liver iron and liver health (Figure 6K and 6L). Additional clinical findings for these cases are provided in Figure 8B. Similarly, we detected phenotypic progression associated with risk alleles, including *AR* STR and *APOE*-ε4 (Figure 6C, 6D, and 6H).

**Figure 6.**
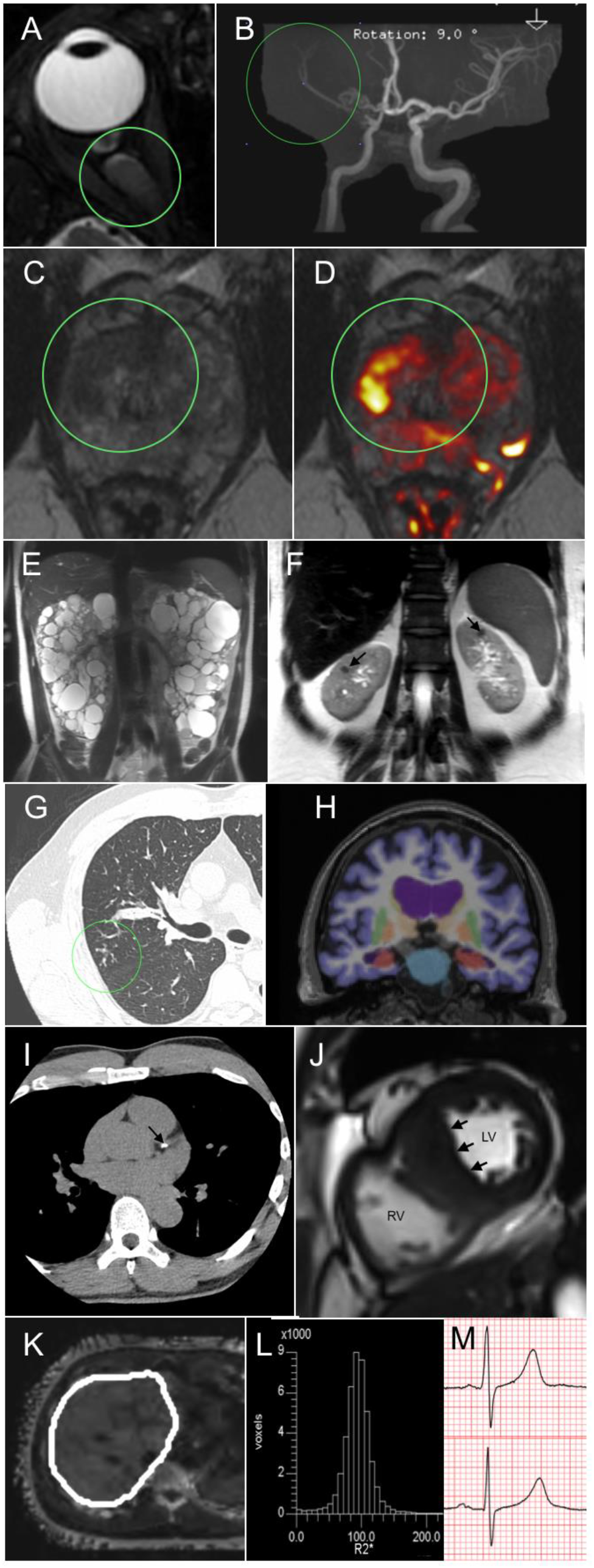
Examples of Integrated Diagnoses. (A&B) A male (20’s) with a clinical diagnosis of neurofibromatosis type 1 (**A**) MRI shows optic nerve glioma, stenosis in the right MCA and (**B**) decreased conspicuity of tertiary MCA branches compared to the left side, compatible with Moyamoya syndrome. (**C&D**) Male with prostate cancer; AR CAG repeats:7; PSA:16.13; Gleason 3+4+5 based on surgical specimen. Green circles mark a suspicious lesion on (**C**) T2 FSE and (**D**) Overlay of color-coded DWI (B1000; NSA=8) onto the T2-weighted image. (**E**) A male (40’s) heterozygous for a likely pathogenic *PKD1* variant, Coronal T2-weighted image shows the extent of the disease. (**F**) A female (40’s) heterozygous for a *HNF1B* likely pathogenic variant with renal cysts and diabetes syndrome. Black arrows on the coronal T2-weighted image point to bilateral sub-centimeter renal cysts with blood products. (**G**) A male (60’s) compound heterozygous for CFTR pathogenic variants with digestive issues/bloating, and chronic sinus infections. CT of the lungs revealed tree in bud nodularity at the right mid and upper lung zone. (**H**) A male (70’s) heterozygous for APOE-ε4 allele; Segmented 3D T 1-weighted MRI of the brain shows enlarged lateral and inferior lateral ventricles along with smaller hippocampi (HOC=0.63, hippocampal volume at 9^th^ percentile compared to age-matched references; assessment with an in-house AI-based brain segmentation algorithm; Wu et al. submitted). (**I**) A male (50’s) heterozygous for a pathogenic variant in the *PCSK9* gene. Black arrow points to the calcified left main/anterior descending coronary artery. (**J**) A male (60’s) heterozygous for a *MYH7* pathogenic variant. MRI shows the hypertrophic ventricular septum consistent with hypertrophic cardiomyopathy (black arrows; LV:left ventricle; RV:right ventricle). (**K&L**) A male (50’s) compound heterozygous for pathogenic variants in the *HFE* gene and has an elevated R2* level (95); (**K**) shows the segmented liver from which R2* data was acquired; (**L**) A histogram of the R2* level by voxel (Siemens liver lab). (**M**) A male (40’s) heterozygous for a likely pathogenic *KCNQ1* variant with a fainting during swimming and multiple events of syncope. ECG showed a prolonged QTc interval (QT/QTc - 490/464; bottom chart compared to someone with a normal QTc interval; top chart).

### Echocardiography, ECG, and computed tomography (CT) scan, and 2-week cardiac rhythm monitoring for cardiovascular disease detection

Among our participants, 16.4% had aberrant cardiac structural or function findings. Left ventricular (LV) hypertrophy was detected in 119 participants with 6 cases considered moderate to severe. The bicuspid aortic valve was observed in eight participants, similar to known prevalence^69^, and we identified a variant (c.1428C>G, p.Tyr476*) in the *SMAD6* gene for one participant with such condition. *MYBPC3* (n=8), *MYH7* (n=5), *MYL2* (n=4) were the most commonly affected genes with relevant cardiac structural findings or family history. Long QT (n=3) and Brugada (n=2) syndromes were the most common affected genetic conditions with related conduction disorders or arrhythmia findings. Details of clinically significant findings on cardiac structural/function, conduction disorders or arrhythmia are listed in Supplemental Table S7A and S7B.

Atherosclerotic cardiovascular disease is the leading cause of death, and coronary artery disease (CAD) accounts for half of all such deaths^70^. Elevated lipids are one of the risk factors for CAD disease. *LDLR* (n=6), *APOB* (n=5), *PCSK9* (n=3), and *LPL* (n=3) were most affected genes associated with familial hypercholesterolemia and hyperlipidemia with correlated clinical presentations. We employed the CT scan for coronary risk stratification. Coronary artery calcium (CAC) scores over 300 were identified together with risks associated with elevated cholesterol (n=66), diabetes (n=104), hypertension (n=30), and chronic renal disease (n=7) (Supplemental Table S8). Two patients had none of the conventional risk factors included in professional guidelines for CAD disease stratification. We identified a LOF variant, c.63dupA (p.Leu22fs) in the *LMBRD1* gene, associated with methylmalonic aciduria and homocystinuria, cblF type (MIM, 277380), in a 61-year old female with a CAC score of 2963. The relative risk of coronary heart disease is estimated as 10.8 for individuals with CAC scores more than 1000^71,72^. However, this individual had a low 10-year risk Framingham score (<5%) which would not have led to a recommendation for a CT scan^73^. The same variant was detected in three relatives of this individual. Two of them had CAC scores of more than 400 and elevated homocysteine levels, suggesting a possible association of homocysteine with vascular calcification. We identified another individual with a variant c.440G>C (p.Gly147Ala) in the *MMACHC* gene, associated with methylmalonic acidemia and homocystinuria (MIM 277400) also had a CAC score of 952. Elevated homocysteine levels have been repeatedly linked to atherosclerosis^74^. A recent publication has shown that homocysteine may contribute to vascular calcification by enhancing osteogenic differentiation of mesenchymal stem cell; thus promoting aortic smooth muscle cell calcification^75,76^. In a case-control study of participants with and without coronary artery calcification (CAC>1), we found an approximately 3-fold increased risk of CAC in those with homocysteine levels higher than 15 mmol/L (OR 2.76, 95% CI [1.4001 to 5.4413]; t-test p=0.03).

### Diabetes Detection

In our cohort, the detection of individuals carrying diabetic risk variants was 0.8% (n=10) which was higher than the known prevalence (0.01-0.05%) for maturity-onset diabetes of the young (MODY). Clinical associations were observed in nine out of ten participants using our full set of screening tests. Figure 7A shows the association of our tests with diabetes detection, and genetic variants were identified in the *MC4R* pathway, and the *HFE* and *CFTR* genes (selected cases). Among our participants, 24.1% had evidence of elevated glucose, and 25% had elevated hemoglobin A1c. Similarly, 34.2% of individuals in our cohort had impaired insulin sensitivity, and 29.5% had impaired glucose sensitivity as measured by metabolomics. Individuals with higher BMI values (>30, 14.8%) had elevated values in IGT, IR, glucose, and A1_c_ assays as demonstrated in Figure 7B. Our results suggest that individualized functional measurements detect type 2 diabetes prior to significant medical consequences and lifestyle adjustments or medications may be helpful for these individuals.

**Figure 7A.**
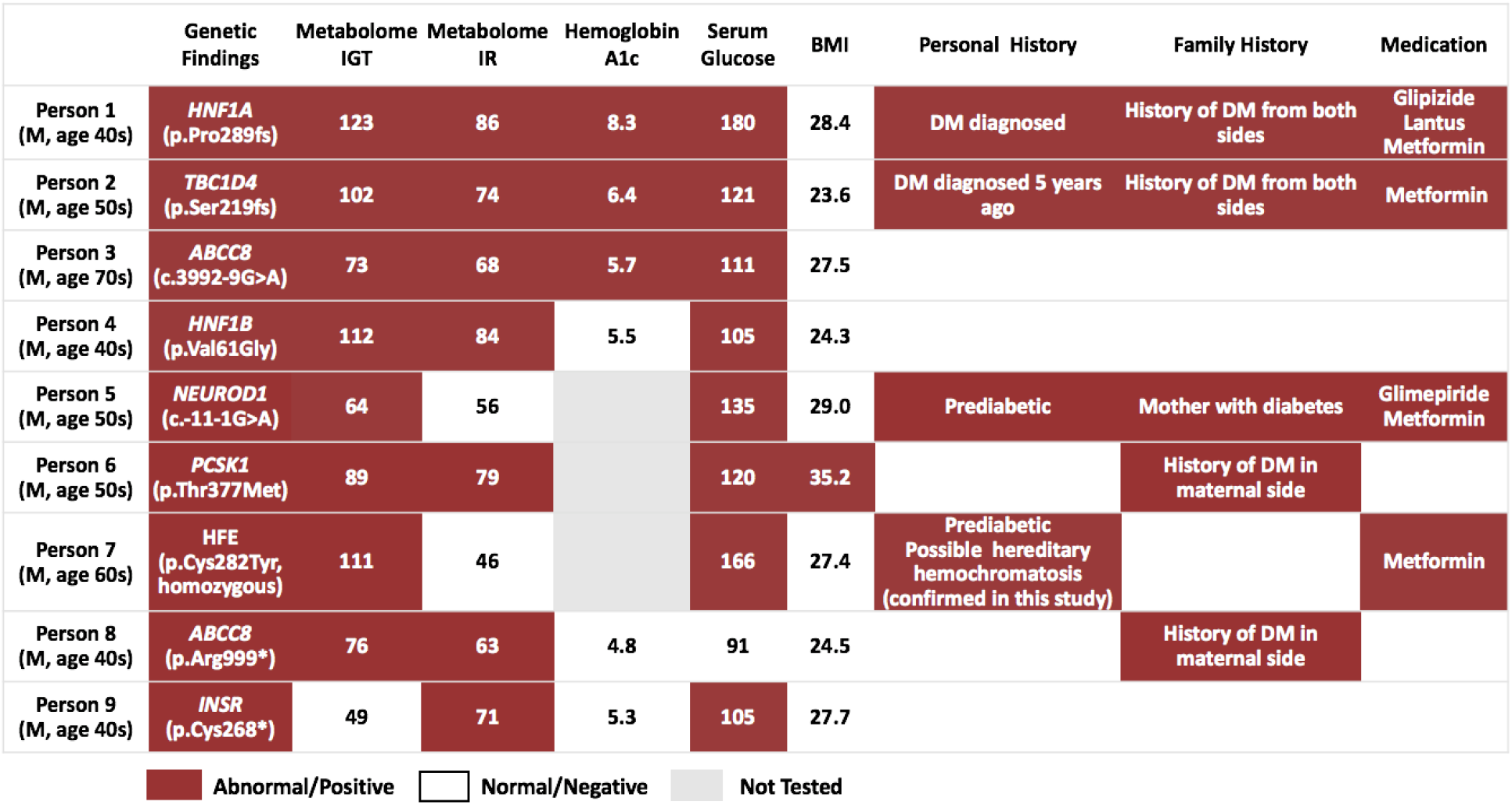
Clinical Associations of Genetic Variants Associated with Diabetes. Nine cases (rows) are presented with nine testing modalities (columns), including genetics, clinical lab measurements, BMI, client and family histories, and current medication. Red cells represent abnormal or positive findings, white cells represent normal or negative findings, and gray cells represent modalities that were not tested. Abnormal/Positive definitions: IGT>60, IR>63, A1c>5.6%, Glucose>99 mg/dL, and BMI>30 kg/m^2^.

**Figure 7B.**
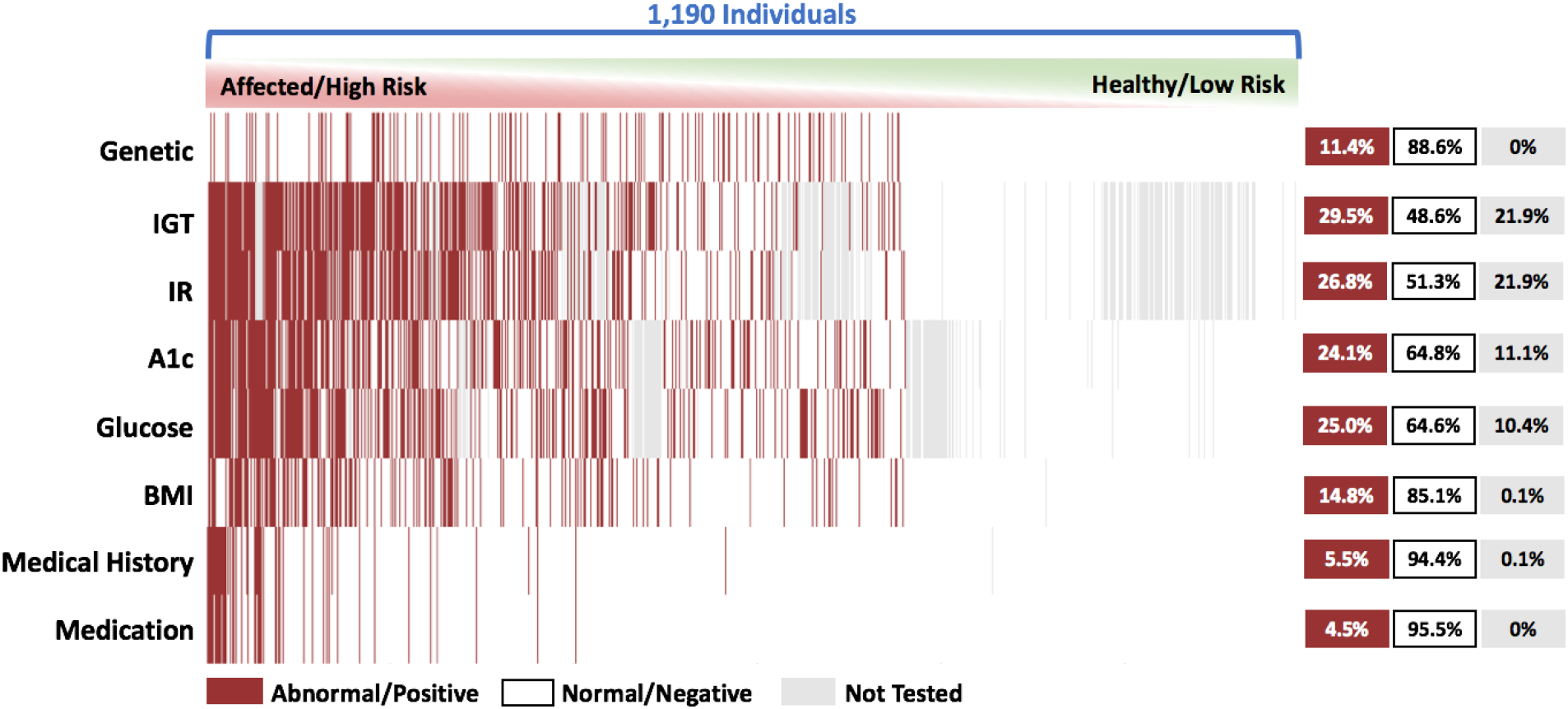
Heatmap Sorted by Number of Abnormal Findings in Diabetic Tests. Red cells represent abnormal or positive findings, white cells represent normal or negative findings, and gray cells represent modalities that were not tested. Abnormal/Positive definitions: IGT>60, IR>63, A1c>5.6%, Glucose>99 mg/dL, and BMI>30 kg/m^2^. The percentage of abnormal/positive findings, normal/negative findings, and not tested findings for each modality is presented to the right.

### Summary of Findings

The summary of results from our data-driven precision health platform is provided in Figure 8A. Details on the selected blood test results, including blood lipids, thyroid function, cancer antigen, liver/kidney function, hematology, hormone, and immunology are listed in Supplemental Table S9. To illustrate the data integration of our tests, we have selected 11 cases (Figure 8B). Case 2 is a participant with no abnormal findings from clinical tests with only a pathogenic variant identified for an autosomal recessive condition. Data obtained in cases 3 to 11 assisted clinicians in newly diagnosing inherited diseases. In particular, case 11 illustrated multiple disease diagnoses. These cases demonstrated that comprehensive testing might allow clinicians to achieve precise clinical assessment and advance disease diagnosis for previously undiagnosed conditions.

**Figure 8A.**
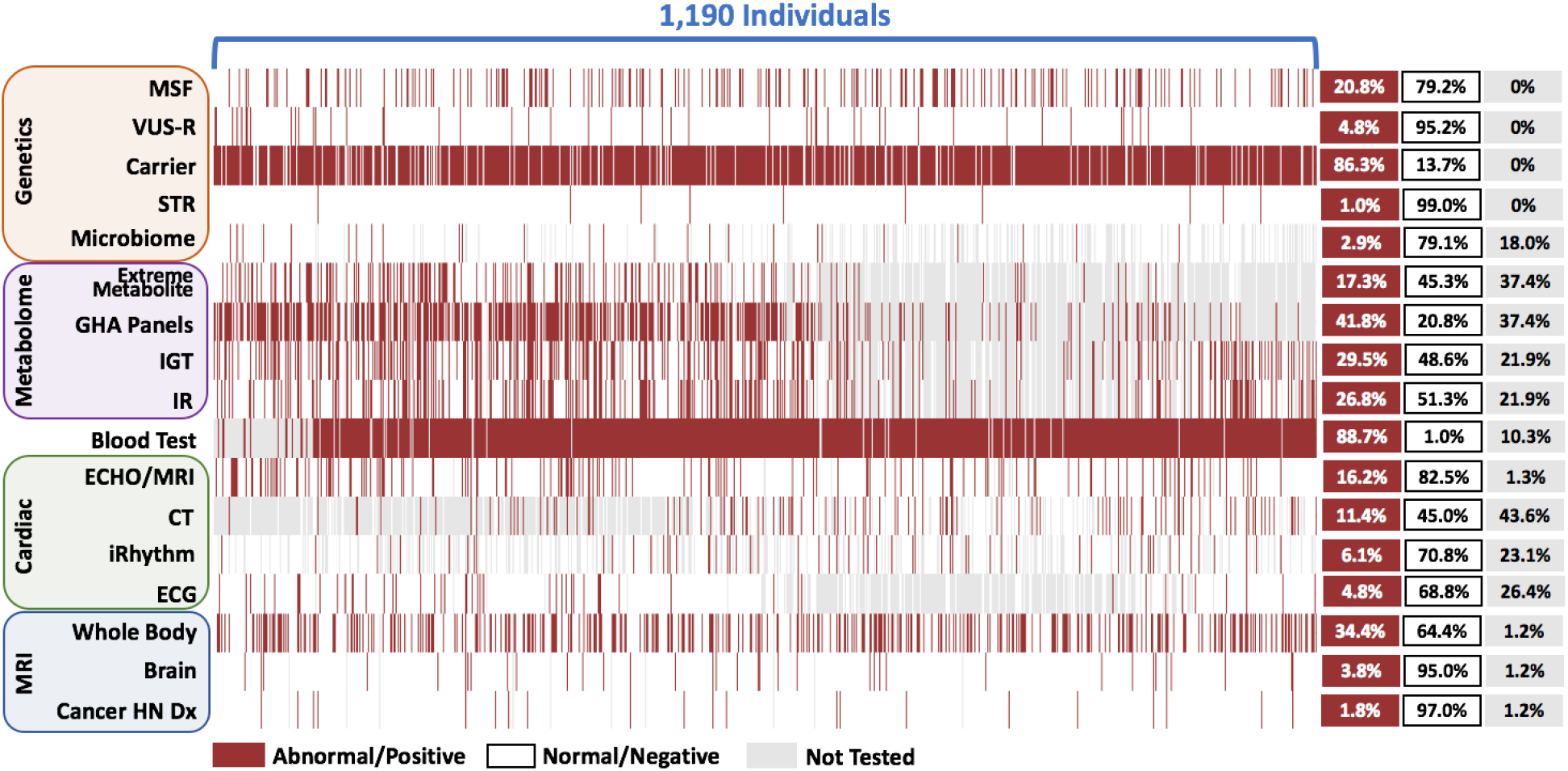
Heatmap of All Test Results (Listed Chronologically). See Methods section in main text for full details. Abbreviations: MSF, medically significant genetic findings; VUS-R, reportable variant of unknown significance; STR, short-tandem repeat; GHA Panels, Global Health Assessment panel; IGT, Quantose impaired glucose tolerance; IR, Quantose insulin resistance; ECHO/MRI, union of dataset of echocardiography and cardiac magnetic resonance imaging; CT, computed tomography coronary artery calcium scoring; ECG, electrocardiogram; Cancer HN Dx, previously undiagnosed cancers.

**Figure 8B.**
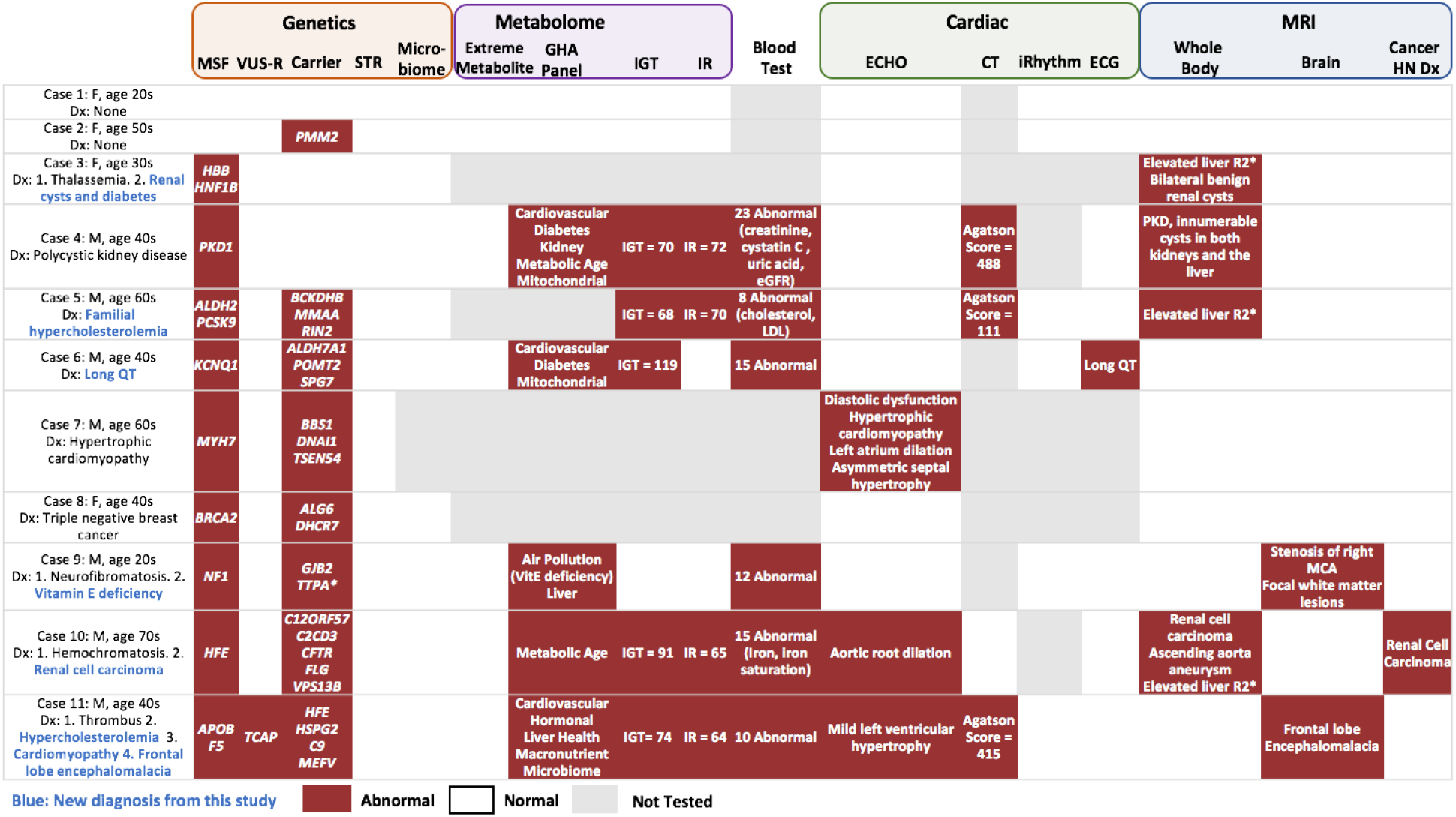
Examples of Integrated Diagnoses. Eleven cases (rows) are presented with 17 testing modalities (columns), including genetics, metabolome, blood test, cardiac and MRI. Red cells represent abnormal or positive findings, white cells represent normal or negative findings, and gray cells represent modalities that were not tested. Abnormal/Positive definitions: IGT>60, IR>63, CT Agatson score>100. See Methods section in main text for full details. Abbreviations: MSF, medically significant genetic findings; VUS-R, reportable variant of unknown significance; STR, short-tandem repeat; GHA Panels, Global Health Assessment panel; IGT, Quantose impaired glucose tolerance; IR, Quantose insulin resistance; ECHO/MRI, union of dataset of echocardiography and cardiac magnetic resonance imaging; CT, computed tomography coronary artery calcium scoring; ECG, electrocardiogram; Cancer HN Dx, previously undiagnosed cancers.

To evaluate the age dependence on our screening tests, we performed an analysis of the age distribution of clinically significant findings (Figure 8C). As expected, the identification of MSF, including carriers of SNVs and those with differences in STR risk, had no age dependency. The metabolite abnormalities were age-dependent in our adult cohort, and the metabolites associated with diabetes contributed a large fraction of the abnormal results. The clinically significant findings detected by ECHO, CT scan, and heart rhythm monitor were age-dependent (p<10^−8^), and the median age of individuals with findings was older than 60 years (ECHO: 62 yrs, CT scan: 65 yrs, heart rhythm: 64 yrs). The clinically significant findings detected by WB-MRI were also age-dependent (p<10^−7^), and in particular, the median age of individuals with findings was 70 years for brain aneurysms/low hippocampal occupancy score and 64.5 years for cancer.

**Figure 8C.**
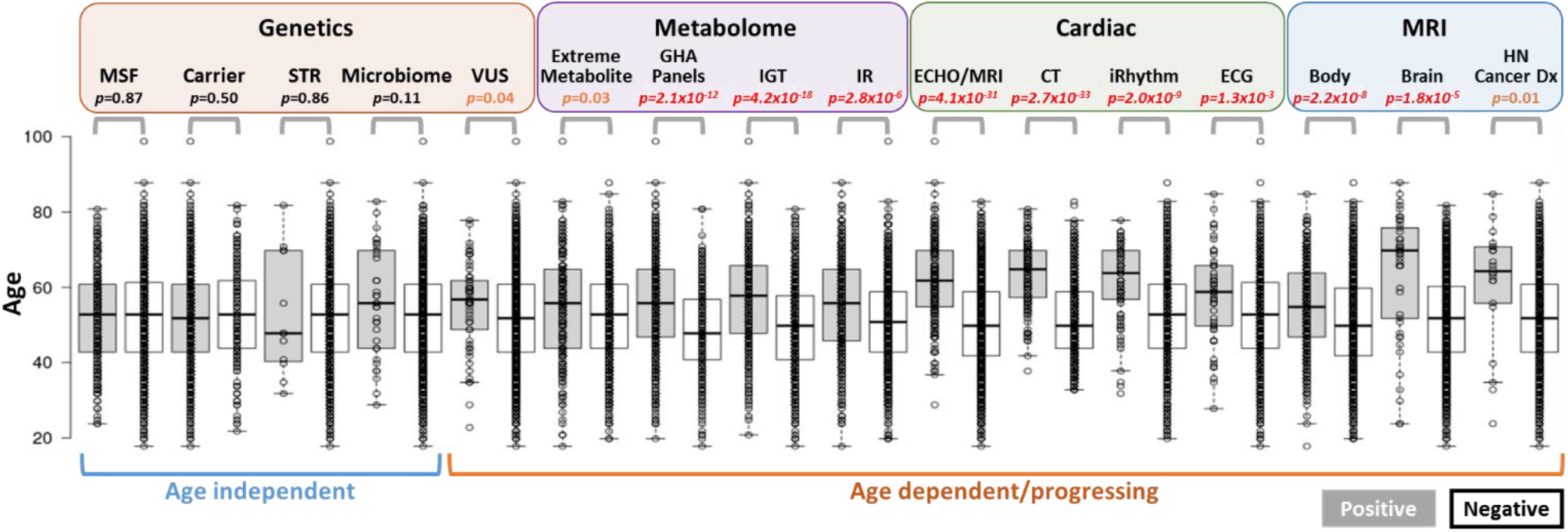
Distribution of Ages in the Cohort with Respect to Positive or Negative Clinically Significant Findings by Test. No statistically significant age dependency (p<0.05) is seen in any clinical significant genetic assessment except for VUS-R. However, all other modalities show age-dependent statistical findings (p<0.05). See Methods section in main text for full details. Abbreviations: MSF, medically significant genetic findings; STR, short-tandem repeat; VUS-R, reportable variant of unknown significance; GHA Panels, Global Health Assessment panel; IGT, Quantose impaired glucose tolerance; IR, Quantose insulin resistance; ECHO/MRI, union of dataset of echocardiography and cardiac magnetic resonance imaging; CT, computed tomography coronary artery calcium scoring; ECG, electrocardiogram; Cancer HN Dx, previously undiagnosed cancers.

## DISCUSSION

Medicine has traditionally been practiced on individuals who are already symptomatic. Clinicians integrate test results for differential diagnosis and devise treatments accordingly. The incorporation of new technologies into medical practice, including DNA sequencing, and non-invasive imaging, enable proactive early detection of disease. Precision medicine initiatives such as MedSeq^77^ and DiscovEHR^53,78^, have reported successes in the early detection of disease when combining conventional clinical practices with genomic sequencing. Despite these successes, the implementation of genomic sequencing in the health care system for healthy individuals for the detection of genetic risk has not yet been accepted^79,80^.

Results from our precision health platform showed that 24.3% of individuals had medically significant genetic findings (MSF) associated with an increased risk of medical disorders, including hematological, cancer, cardiovascular, endocrine, and neurological. A total of 206 unique medically significant variants in 111 genes were identified. Our findings showed that 63.1% of individuals with MSF had associated phenotypes observed in phenotypic testing, enabling clinicians to integrate genetic analysis into their assessments for clinical-molecular diagnosis. Among disease categories, we found that cardiovascular and endocrine diseases achieved considerable clinical associations between MSF and clinical phenotypes (89% and 72% respectively). The probability of a non-pathogenic MSF (1% to 10% for a likely pathogenic variant), reduced penetrance, or late-onset of disease presentation may be reasons for the remaining lack of associations. The detection of MSF enables clinical interventions to improve future health outcomes, the optimal use of treatment prior to the deterioration of the condition and the early detection of disease for asymptomatic family members.

A precise clinical-molecular diagnosis can provide individually tailored treatment or surveillance options. For example, the clinical-molecular diagnosis of maturity onset of diabetes of the young can provide precise treatment such as low-dose sulfonylureas for affected individuals without a possible incorrect use of insulin as the initial treatment ^81^. The consensus guideline published by The Familial Hypercholesterolemia (FH) Foundation estimates that many FH patients remain undiagnosed and recommend FH genetic testing to improve diagnosis, initiate therapies at an earlier age, and provide prognostic assessment^82^. It has been estimated that ~1% of the U.S. population has a plausible genetic risk for cancer or cardiovascular disease^80^. Thus, early detection of these at-risk individuals via a genomic approach seems to be a reasonable goal to mitigate the risk effectively and prevent premature mortality. We found that ~75% (39/51) of participants with a P/LP cancer gene variant did not meet pertinent genetic testing criteria (unpublished results), suggesting current genetic testing guidelines may be inadequate to identify at-risk individuals. There is a demanding need for extensive studies that facilitate refinement of risk estimates and facilitate the implementation of DNA analysis as genomic screening for routine clinical care.

From the Institute of Health Metrics and Evaluation, age-related chronic disease is the leading cause of pre-mature mortality in U.S. adults between 50-74 years of age, affecting 39% of men and 24% of women^81^. Our precision health platform identified clinically significant findings in disease groups associated with premature mortality, including cancer, cardiovascular, endocrine, cirrhosis, and neurologic conditions. In our presumed healthy participants, we identified 19 individuals (2%) with early-stage neoplasia, prostate adenocarcinoma, renal cell carcinoma, lymphoma, transitional cell carcinoma, papillary thyroid cancer, pancreatic cancer and mediastinal thymoma that required prompt (<30 days) medical attention. Additionally, we identified 29 individuals (2.4%) with severe cardiovascular clinically significant findings that required prompt medical attention: 18 participants with an enlarged aortic root, 8 with newly recognized atrial fibrillation, 2 with moderate-to-severe left ventricular hypertrophy, and 1 with severe asymmetric septal hypertrophy consistent with hypertrophic cardiomyopathy. These results demonstrate that our precision health platform enable the early detection of at-risk individuals; thus, preventative measures can be employed to reduce premature mortality.

Individualized (N-of-1) findings from our integrated precision health platform include 1. Detection of medically significant genetic findings (MSF) which contributes to an increased risk of disease; 2. Detection of quantitative imaging biomarkers that allow the assessment of individualized health status; 3. Early detection of diabetes that allows lifestyle modifications or preventative measures for the risk mitigation; 4. Early detection of individuals with a high cardiovascular risk that allows the use of preventative measures; 5. Personalized drug response measurement or supplement uses by metabolomics, and others yet to be identified. We will evaluate these findings in larger numbers of participants over extended periods of time. Test results from repeat evaluations will provide baselines per individual and may improve our ability to diagnosis and prevent or manage chronic diseases.

Our findings associate a lower number of CAG repeats within the *AR* gene with prostate growth and/or cancer risk. These lower number of CAG repeats have been previously associated with elevated *AR* expression. This finding is of therapeutic interest, given the availability of 5-alpha-hydrogenase inhibitors which have been shown to decrease *AR* expression^83^. The 5-alpha-hydrogenase inhibitors have been reported to lower the incidence of prostate cancer and are an FDA-approved treatment for benign prostatic hyperplasia (BPH)^84^. The association of the *AR* CAG repeats with prostate cancer and BPH have been elusive and controversial^60,61,62^, and we will continuously monitor the progression of prostate diseases with iterative clinical evaluation. African American individuals who have a higher risk of prostate cancer as indicated by the National Comprehensive Cancer Network guidelines and have short STR in the *AR* gene would be a logical cohort for longitudinal studies (Figure 4). The combination of non-contrast MR imaging and genetic testing may have the potential to provide individualized decision support for BPH and prostate cancer.

Our study shows that with in-depth quantitative testing, individuals who are carriers for autosomal recessive diseases exhibit detectable phenotypic changes given the sensitivity of WB-MRI and metabolomics. We observed that some carriers of known autosomal recessive conditions had a penetrance for metabolomics biomarkers and imaging abnormalities. The *PKHD1* carriers had numerous liver and/or kidney cysts detected by MRI and repeat evaluation by imaging allows clinicians to determine whether these individuals may have isolated polycystic liver disease. *HFE* carriers (p.Cys282Tyr) had elevated R2* detected by MRI, suggesting the regulation of iron is compromised. Symptoms associated with hereditary hemochromatosis may need to be evaluated in these individuals over longer periods. For metabolic penetrance, 10/30 (33%) *PAH* carriers had moderately elevated phenylalanine detected by metabolomics testing. Thirty-four (13%) carriers of pathogenic variants identified by WGS in the genes associated with metabolism pathways had elevated or decreased (95% CI) levels in the corresponding metabolites. These data are not explored extensively to evaluate if long-term penetrance may cause a clinical impact on health. Repeat evaluation of these individuals is required to characterize the clinical significance of undefined findings.

The precision medicine initiative has the potential to change the practice of medicine from reactive to proactive and preventative via early detection and risk mitigation. Our data achieved some notable near-term successes and support a research and clinical emphasis beyond genome sequencing.

## AUTHOR CONTRIBUTIONS

J.C.V. conceived the study. C.T.C. led the analyses. Y,-C.C.H., H.-C.Y., R.M., and C.T.C. performed genomic analyses and genetic-phenotypic integration analysis. N.M.S.-A., C.L.S., S.D., N.H., A.M.K., and D.S.K. evaluated clinical imaging. Y,-C.C.H., H.-C.Y., R.M., R.H., J.B., C.R., E.S., K.D., M.D., P.B., and C.T.C. contributed to clinical genetics evaluation. Y,-C.C.H., H.-C.Y., E.T.C., I.V.C., and T.J.J. performed metabolomics analysis. L.N. contributed to data collection. H.T., W.L., M.H., and E.F.K. contributed to bioinformatics. R.M. R.H., J.B., C.R., E.S., K.D., M.D., P.B., A.M.K., D.S.K., and C.T.C. contributed to clinical support. D.S.K. and C.T.C supervised research. Y,-C.C.H., H.-C.Y., N.M.S.-A., and C.T.C. wrote the paper. All authors reviewed and approved the manuscript.

## Supporting information

## ACKNOWLEDGMENTS

The authors would like to acknowledge the individuals who participated in this novel precision medicine study without whom the findings would not be possible. Julie Ellison provided medical editing assistance. The authors would like to acknowledge the Human Longevity and Health Nucleus staff past and present: Amy Reed, Alexander Graff, Ana Sanchez, Athena Hutchinson, Brad Perkins, Carina Sarabia, Cheryl Buffington, Cheryl Greenberg, Christina Bonas, Catherine Vrona, Daniel Jones, Danielle Dorris, Desiree Cheney, Dmitry Tkach, Diana Cardin Escobedo, Eric Dec, Genelle Olsen, Greg Olson, Heidi Millard, Helen Messier, Hyun-Kyung Chung, Jason Deckman, Javier Velazquez-Muriel, Jian Wu, Kathy Levine, Keisha Robinson, Krista-Lynn Banner, Kristi Ericksen-Miller, Laura Edwards, Melissa Schweitzer, Nicole Boramanand, Nolan Tengonciang, Patrick Jamieson, Samantha Punsalan, Tom Folan, Victor Lavrenko, Wayne Delport, and William Herrera.

