## Supplementary material for "Precision Medicine Advancements Using Whole Genome Sequencing, Noninvasive Whole Body Imaging, and Functional Diagnostics"

**Supplemental Table S1. Non-contrast Whole-body Magnetic Resonance Imaging.**

| Region of Interest | Protocol | Purpose |
|---|---|---|
| **Brain** | 3D T1 MPRAGE/FSPGR | Structural analysis/segmentation. Post-processing with Neuroquant® prospectively; post-processing with in-house AI-based algorithm retrospectively |
| | 3D FLAIR | White matter hyperintensity and gliosis assessment |
| | RSI (B500, 1500, 4000; 6, 6, and 12 directions with 2 B0 directions) or 2D DWI (B1000 NSA=2, B0) | Evaluation of tumors and hypercellular pathology |
| | T2* 2D GRE | Calcification and blood product detection (subset of cohort) |
| | 3D Time-of-Flight MRA | Cerebrovascular disease |
| **Whole body (five stations: head, neck, chest, abdomen, pelvis)** | T1 Dixon imaging | Anatomic overview and AMRA post-processing for body composition (subset of cohort) |
| | T2 SSFSE/HASTE | Anatomic overview |
| | DWI (B1000 NSA=5, B0) | Primarily for identification of tumors |
| **Cardiac** | 2,3,4 CH LAX CINE, SAX stacked cine imaging | Cardiac structure and function (subset of cohort) |
| **Liver** | Coronal HASTE/SSFSE | Biliary tree |
| | Multi-echo Dixon | Liver fat and iron evaluation |
| **Prostate** | Focused FOV T2 TSE/FSE | Lesion localization |
| | RSI (B125, 375, 1000; 6, 6, and 12 directions with 2 B0 directions or DWI (B1000 NSA=8, B0) | Modified PIRADS evaluation for cancer detection |

DWI: diffusion weighted imaging; FLAIR: fluid-attenuated inversion recovery; FSE: fast spin echo; FSPGR: fast spoiled gradient echo; GRE: gradient echo; HASTE: half Fourier single-shot turbo spin echo; MPRAGE: magnetization-prepared 180 degrees radio-frequency pulses and rapid gradient-echo; MRA: magnetic resonance angiography; NSA: number of signal averages; PIRADS: prostate imaging reporting and data system; RSI: restriction spectrum imaging; SSFSE: single shot fast spin echo; TSE: turbo spin echo.

**Supplemental Table S2. Cardiovascular Abnormal Criteria.**

| **Abnormal Cardiac Rhythm Criteria** |
|---|
| Atrial fibrillation or atrial flutter |
| Sustained ventricular tachycardia (>30 seconds) |
| Atrioventricular block (2$^{nd}$ type II, high degree, or 3$^{rd}$ degree) |
| Sinus pauses lasting more than 3 seconds |
| Bundle branch blocks (QRS >0.120 mg): Left, Right, Nonspecific |
| Abnormal QTc interval: elongated => 450 ms (men), => 460 ms (women) |
| Pre-excitation (Wolff-Parkinson-White) (>1% premature atrial contractions; >5% premature ventricular contractions) |
| Supraventricular tachycardia lasting more than 20 beats in any single run |

| **Echocardiogram Abnormal Findings Criteria** | |
|---|---|
| **Structure of Interest** | **Abnormal findings criteria** |
| **Left ventricle** | - any systolic dysfunction or wall motion abnormality<br>- moderate or severe diastolic dysfunction<br>- mild or worse hypertrophy<br>- dynamic outflow obstruction<br>- hypertrophic cardiomyopathy |
| **Right ventricle** | - systolic dysfunction<br>- moderate or severe dilation |
| **Left or Right Atrium** | - moderate or severe dilation |
| **Valves** | - aortic valve: regurgitation that is mild or worse<br>- Tricuspid, pulmonary and mitral valves: regurgitation that is mild-moderate or greater<br>- valvular stenosis<br>- bicuspid aortic valve<br>- mitral valve prolapse |
| **Pericardium** | - pericardial effusion (small or larger) |
| **Pulmonary artery pressure** | - mild or worse elevation of pressure artery pressure |
| **Abnormal cardiac echodensity/mass** | - evidence of mass/tumor/thrombus |
| **Atrial or ventricular septum** | - structural defects |
| **Aortic root** | - dilation with diameter => 4.4cm (males) or 4.0cm (females) |

### Supplemental Table S3: Case Details and Corresponding Clinical Correlations Detected in Health Nucleus Screening Tests

| Variant | Gene | Disease | MOI | ACMG Classification | Variant Class | Zygosity | Phenotype confirmation | Family History Confirmation |
|---|---|---|---|---|---|---|---|---|
| chr11:17397055 C>T \| c.3992-9G>A | ABCC8 | Familial hyperinsulinemic hypoglycemia | AD/AR | P | MSF | Heterozygous | Y (Blood test, Metabolome) | N |
| chr12:111803962 G>A \| p.Glu504Lys \| c.1510G>A | ALDH2 | Alcohol sensitivity | AD | P | MSF | Heterozygous | N | N |
| chr12:111803962 G>A \| p.Glu504Lys \| c.1510G>A | ALDH2 | Alcohol sensitivity | AD | P | MSF | Heterozygous | N | N |
| chr12:111803962 G>A \| p.Glu504Lys \| c.1510G>A | ALDH2 | Alcohol sensitivity | AD | P | MSF | Heterozygous | N | N |
| chr12:111803962 G>A \| p.Glu504Lys \| c.1510G>A | ALDH2 | Alcohol sensitivity | AD | P | MSF | Heterozygous | N | Y |
| chr12:111803962 G>A \| p.Glu504Lys \| c.1510G>A | ALDH2 | Alcohol sensitivity | AD | P | MSF | Heterozygous | N | N |
| chr12:111803962 G>A \| p.Glu504Lys \| c.1510G>A | ALDH2 | Alcohol sensitivity | AD | P | MSF | Homozygous | N | N |
| chr12:111803962 G>A \| p.Glu504Lys \| c.1510G>A | ALDH2 | Alcohol sensitivity | AD | P | MSF | Heterozygous | N | N |
| chr12:111803962 G>A \| p.Glu504Lys \| c.1510G>A | ALDH2 | Alcohol sensitivity | AD | P | MSF | Heterozygous | N | N |
| chr12:111803962 G>A \| p.Glu504Lys \| c.1510G>A | ALDH2 | Alcohol sensitivity | AD | P | MSF | Heterozygous | N | N |
| chr12:111803962 G>A \| p.Glu504Lys \| c.1510G>A | ALDH2 | Alcohol sensitivity | AD | P | MSF | Heterozygous | N | N |
| chr12:111803962 G>A \| p.Glu504Lys \| c.1510G>A | ALDH2 | Alcohol sensitivity | AD | P | MSF | Heterozygous | N | N |
| chr12:111803962 G>A \| p.Glu504Lys \| c.1510G>A | ALDH2 | Alcohol sensitivity | AD | P | MSF | Heterozygous | N | N |
| chr12:111803962 G>A \| p.Glu504Lys \| c.1510G>A | ALDH2 | Alcohol sensitivity | AD | P | MSF | Heterozygous | N | N |
| chr12:111803962 G>A \| p.Glu504Lys \| c.1510G>A | ALDH2 | Alcohol sensitivity | AD | P | MSF | Heterozygous | N | Y |
| chr12:111803962 G>A \| p.Glu504Lys \| c.1510G>A | ALDH2 | Alcohol sensitivity | AD | P | MSF | Heterozygous | N | N |
| chr12:111803962 G>A \| p.Glu504Lys \| c.1510G>A | ALDH2 | Alcohol sensitivity | AD | P | MSF | Heterozygous | N | Y |
| chr12:111803962 G>A \| p.Glu504Lys \| c.1510G>A | ALDH2 | Alcohol sensitivity | AD | P | MSF | Heterozygous | N | N |
| chr12:111803962 G>A \| p.Glu504Lys \| c.1510G>A | ALDH2 | Alcohol sensitivity | AD | P | MSF | Homozygous | N | N |
| chr12:111803962 G>A \| p.Glu504Lys \| c.1510G>A | ALDH2 | Alcohol sensitivity | AD | P | MSF | Homozygous | N | Y |
| chr12:111803962 G>A \| p.Glu504Lys \| c.1510G>A | ALDH2 | Alcohol sensitivity | AD | P | MSF | Homozygous | N | N |
| chr12:111803962 G>A \| p.Glu504Lys \| c.1510G>A | ALDH2 | Alcohol sensitivity | AD | P | MSF | Homozygous | N | Y |
| chr12:111803962 G>A \| p.Glu504Lys \| c.1510G>A | ALDH2 | Alcohol sensitivity | AD | P | MSF | Heterozygous | N | N |
| chr12:111803962 G>A \| p.Glu504Lys \| c.1510G>A | ALDH2 | Alcohol sensitivity | AD | P | MSF | Heterozygous | N | N |
| chr12:111803962 G>A \| p.Glu504Lys \| c.1510G>A | ALDH2 | Alcohol sensitivity | AD | P | MSF | Heterozygous | N | N |
| chr12:111803962 G>A \| p.Glu504Lys \| c.1510G>A | ALDH2 | Alcohol sensitivity | AD | P | MSF | Heterozygous | N | N |
| chr12:111803962 G>A \| p.Glu504Lys \| c.1510G>A | ALDH2 | Alcohol sensitivity | AD | P | MSF | Heterozygous | N | Y |
| chr12:111803962 G>A \| p.Glu504Lys \| c.1510G>A | ALDH2 | Alcohol sensitivity | AD | P | MSF | Heterozygous | N | N |
| chr12:111803962 G>A \| p.Glu504Lys \| c.1510G>A; chr14:23419259 A>T \| p.Leu1297Gln \| c.3890T>A | ALDH2; MYH7 | Alcohol sensitivity; Cardiomyopathy | AD; AD | P; VUS | MSF; VUS-R | Heterozygous; Heterozygous | N; Y (ECHO, ECG) | N; N |

| | | | | | | | | |
|---|---|---|---|---|---|---|---|---|
| chr12:111803962 G>A \| p.Glu504Lys \| c.1510G>A; chr8:54629679 C>T \| p.Arg1933* \| c.5797C>T | ALDH2; RP1 | Alcohol sensitivity; Retinitis pigmentosa | AD; AD/AR | P; VUS | MSF; VUS-R | Heterozygous; Heterozygous | N; Y (PHx) | N; Y |
| chr19:8366271 G>A \| p.Glu167Lys \| c.499G>A | ANGPTL4 | Lower plasma triglyceride level | AD | VUS | VUS-R | Heterozygous | Y (Blood test) | N |
| chr4:113363406 C>T \| p.Leu3609Phe \| c.10825C>T; chr10:74101021 CG>C \| p.Val650fs \| c.1948delG | ANK2; VCL | Cardiac arrhythmia; Cardiomyopathy | AD; AD | VUS; VUS | VUS-R; VUS-R | Heterozygous; Heterozygous | Y (ECHO, ECG); Y (ECHO) | Y; Y |
| chr5:112842768 C>A \| p.Pro2392Thr \| c.7174C>A | APC | Hereditary cancer-predisposing syndrome | AD | VUS | VUS-R | Heterozygous | N | Y |
| chr2:21006288 C>T \| p.Arg3527Gln \| c.10580G>A | APOB | Familial hypercholesterolemia | AD | P | MSF | Heterozygous | Y (Blood test) | Y |
| chr2:21001939 ACTG>A \| p.Gln4494del \| c.13480_13482delCAG | APOB | Familial hypercholesterolemia | AD | VUS | VUS-R | Heterozygous | Y (Blood test) | N |
| chr2:21007956 T>G \| p.Asn2971Thr \| c.8912A>C | APOB | Familial hypercholesterolemia | AD | VUS | VUS-R | Heterozygous | Y (Blood test) | N |
| chr2:21001939 ACTG>A \| p.Gln4494del \| c.13480_13482delCAG | APOB | Familial hypercholesterolemia | AD | VUS | VUS-R | Heterozygous | Y (Blood test) | Y |
| chr11:116830637 C>T \| p.Arg37* \| c.109C>T | APOC3 | Apolipoprotein C-III deficiency | AD | P | MSF | Heterozygous | Y (Blood test) | N |
| chr11:116830637 C>T \| p.Arg37* \| c.109C>T | APOC3 | Apolipoprotein C-III deficiency | AD | P | MSF | Heterozygous | Y (Blood test) | N |
| chr11:108272531 G>A \| c.3078-1G>A | ATM | Hereditary cancer-predisposing syndrom | AD | P | MSF | Heterozygous | N | Y |
| chr11:108250752 CTG>C \| p.Cys430fs \| c.1290_1291delTG | ATM | Hereditary cancer-predisposing syndrom | AD | P | MSF | Heterozygous | N | Y |
| chr11:108316015 C>T \| p.Arg2034* \| c.6100C>T | ATM | Hereditary cancer-predisposing syndrom | AD | P | MSF | Heterozygous | Y (PHx) | Y |
| chr11:108250866 CAA>C \| p.Lys468fs \| c.1402_1403delAA | ATM | Hereditary cancer-predisposing syndrom | AD | P | MSF | Heterozygous | N | Y |
| chr2:214809498 G>GGACGCGGAACGAGGCTCGTTCCC \| p.Ala25fs \| c.71_72insGGGAACGAGCCTCGTTCCGCGTC | BARD1 | Cancer | AD | LP | MSF | Heterozygous | N | Y |
| chr17:43124027 ACT>A \| p.Glu23fs \| c.68_69delAG | BRCA1 | Breast-ovarian cancer | AD | P | MSF | Heterozygous | N | Y |
| chr13:32380043 C>T \| p.Arg3052Trp \| c.9154C>T | BRCA2 | Breast-ovarian cancer | AD | P | MSF | Heterozygous | Y (PHx) | Y |
| chr13:32340072 ACT>A \| p.Leu1908fs \| c.5722_5723delCT | BRCA2 | Breast-ovarian cancer | AD | P | MSF | Heterozygous | N | Y |
| chr13:32329467 CTG>C \| p.Val220fs \| c.658_659delGT | BRCA2 | Breast-ovarian cancer | AD | P | MSF | Heterozygous | N | Y |
| chr13:32337160 TAAAC>T \| p.Ala938fs \| c.2808_2811delACAA | BRCA2 | Breast-ovarian cancer | AD | P | MSF | Heterozygous | N | Y |
| chr13:32326085 CTTTGT>C \| c.426-12_426-8delGTTTT | BRCA2 | Breast-ovarian cancer | AD | VUS | VUS-R | Heterozygous | N | Y |
| chr13:32339143 TTC>T \| p.Leu1598fs \| c.4793_4794delTC | BRCA2 | Breast-ovarian cancer | AD | LP | MSF | Heterozygous | N | N |
| chr13:32329467 CTG>C \| p.Val220fs \| c.658_659delGT | BRCA2 | Breast-ovarian cancer | AD | P | MSF | Heterozygous | N | N |
| chr17:61716051 G>A \| p.Arg798* \| c.2392C>T | BRIP1 | Cancer-predisposing syndrome | AD | P | MSF | Heterozygous | N | N |
| chr17:61716043 G>C \| p.Tyr800* \| c.2400C>G | BRIP1 | Cancer-predisposing syndrome | AD | P | MSF | Heterozygous | N | Y |
| chr17:61685976 A>C \| p.Leu922* \| c.2765T>G; chr11:46739505 G>A \| c.*97G>A | BRIP1; F2 | Cancer-predisposing syndrome; Thrombophilia | AD; AD | P; P | MSF; MSF | Heterozygous; Heterozygous | N; N | N; N |

| | | | | | | | | |
|---|---|---|---|---|---|---|---|---|
| chr3:15645186 G>C \| p.Asp446His \| c.1336G>C | BTD | Biotinidase deficiency | AR | LP | MSF | Homozygous | N | N |
| chr3:15645186 G>C \| p.Asp446His \| c.1336G>C | BTD | Biotinidase deficiency | AR | LP | MSF | Homozygous | N | N |
| chr3:15644367 G>A \| p.Ala173Thr \| c.517G>A; chr3:15645186 G>C \| p.Asp446His \| c.1336G>C | BTD; BTD | Biotinidase deficiency; Biotinidase deficiency | AR; AR | P; LP | MSF; MSF | Heterozygous; Homozygous | N; N | N; N |
| chr3:15645063 T>G \| p.Phe405Val \| c.1213T>G; chr3:15645186 G>C \| p.Asp446His \| c.1336G>C | BTD; BTD | Biotinidase deficiency; Biotinidase deficiency | AR; AR | P; LP | MSF; MSF | Heterozygous; Heterozygous | N; N | N; N |
| chr1:196994128 C>CA \| p.Glu163fs \| c.486dupA | CFHR5 | Nephropathy due to CFHR5 deficiency | AD/AR | VUS | VUS-R | Heterozygous | Y (PHx) | N |
| chr22:28695219 G>A \| p.Ser471Phe \| c.1412C>T | CHEK2 | Hereditary cancer-predisposing syndrome | AD | P | MSF | Heterozygous | N | Y |
| chr22:28734532 C>T \| p.Glu64Lys \| c.190G>A | CHEK2 | Hereditary cancer-predisposing syndrome | AD | VUS | VUS-R | Heterozygous | N | Y |
| chr22:28695868 AG>A \| p.Thr410fs \| c.1229delC | CHEK2 | Hereditary cancer-predisposing syndrome | AD | P | MSF | Heterozygous | N | N |
| chr22:28695219 G>A \| p.Ser471Phe \| c.1412C>T | CHEK2 | Hereditary cancer-predisposing syndrome | AD | P | MSF | Heterozygous | N | N |
| chr22:28725099 A>G \| p.Ile200Thr \| c.599T>C | CHEK2 | Hereditary cancer-predisposing syndrome | AD | LP | MSF | Heterozygous | N | Y |
| chr22:28725099 A>G \| p.Ile200Thr \| c.599T>C | CHEK2 | Hereditary cancer-predisposing syndrome | AD | LP | MSF | Heterozygous | N | N |
| chr22:28725099 A>G \| p.Ile200Thr \| c.599T>C | CHEK2 | Hereditary cancer-predisposing syndrome | AD | LP | MSF | Heterozygous | N | N |
| chr22:28725099 A>G \| p.Ile200Thr \| c.599T>C | CHEK2 | Hereditary cancer-predisposing syndrome | AD | LP | MSF | Heterozygous | N | N |
| chr22:28695868 AG>A \| p.Thr410fs \| c.1229delC | CHEK2 | Hereditary cancer-predisposing syndrome | AD | P | MSF | Heterozygous | Y (PHx) | Y |
| chr22:28725099 A>G \| p.Ile200Thr \| c.599T>C | CHEK2 | Hereditary cancer-predisposing syndrome | AD | LP | MSF | Heterozygous | N | N |
| chr22:28734532 C>T \| p.Glu64Lys \| c.190G>A | CHEK2 | Hereditary cancer-predisposing syndrome | AD | VUS | VUS-R | Heterozygous | N | Y |
| chr6:70281388 A>T \| c.876+2T>A | COL9A1 | Stickler syndrome | AD | LP | MSF | Heterozygous | Y (PHx) | Y |
| chr3:98591046 CATTTG>C \| p.Gln221fs \| c.661_665delCAAAT | CPOX | Coproporphyria | AD | LP | MSF | Heterozygous | N | N |
| chr16:50792641 CT>C \| p.Phe763fs \| c.2289delT; chr19:11133635 C>G \| c.*2117C>G | CYLD; LDLR | Cylindromatosis/Trichoepithelioma; Familial hypercholesterolemia | AD; AD | LP; VUS | MSF; VUS-R | Heterozygous; Heterozygous | Y (PHx); Y (Blood test) | Y; Y |
| chr6:32040110 G>T \| p.Val282Leu \| c.844G>T | CYP21A2 | Adrenal hyperplasia | AR | LP | MSF | Heterozygous | N | N |
| chr6:32040110 G>T \| p.Val282Leu \| c.844G>T | CYP21A2 | Adrenal hyperplasia | AR | LP | MSF | Heterozygous | N | N |
| chr5:79029971 C>A \| p.Val583Phe \| c.1747G>T; chr5:79033302 T>A \| p.Lys434* \| c.1300A>T | DMGDH; DMGDH | Dimethylglycine dehydrogenase deficiency; Dimethylglycine dehydrogenase deficiency | AR; AR | VUS; LP | MSF; MSF | Heterozygous; Heterozygous | Y (Metabolome); Y (Metabolome) | N; N |
| chr18:31074864 G>T \| p.Asp569Glu \| c.1707C>A | DSC2 | Arrhythmogenic right ventricular cardiomyopathy | AD | VUS | VUS-R | Heterozygous | N | Y |
| chr18:31091123 C>A \| p.Glu127* \| c.379G>T | DSC2 | Arrhythmogenic right ventricular cardiomyopathy | AD | LP | MSF | Heterozygous | N | N |
| chr1:231421829 G>C \| p.Tyr20* \| c.60C>G | EGLN1 | Erythrocytosis | AD | LP | MSF | Heterozygous | Y (Blood test) | N |
| chr9:127819966 C>CTT \| p.Phe403fs \| c.1204_1205dupAA | ENG | Hereditary hemorrhagic telangiectasia | AD | P | MSF | Heterozygous | Y (PHx) | Y |
| chr2:47375300 G>A \| c.575+1G>A | EPCAM | Lynch syndrome | AD | LP | MSF | Heterozygous | N | Y |
| chr4:186280258 T>C \| p.Phe301Leu \| c.901T>C | F11 | Factor XI deficiency | AD/AR | P | MSF | Heterozygous | N | N |

| | | | | | | | | |
|---|---|---|---|---|---|---|---|---|
| chr11:46739505 G>A \| c.*97G>A | F2 | Thrombophilia | AD | P | MSF | Heterozygous | N | N |
| chr11:46739505 G>A \| c.*97G>A | F2 | Thrombophilia | AD | P | MSF | Heterozygous | N | N |
| chr11:46739505 G>A \| c.*97G>A | F2 | Thrombophilia | AD | P | MSF | Heterozygous | N | N |
| chr11:46739505 G>A \| c.*97G>A | F2 | Thrombophilia | AD | P | MSF | Heterozygous | N | N |
| chr11:46739505 G>A \| c.*97G>A | F2 | Thrombophilia | AD | P | MSF | Heterozygous | N | N |
| chr11:46739505 G>A \| c.*97G>A | F2 | Thrombophilia | AD | P | MSF | Heterozygous | N | N |
| chr11:46739505 G>A \| c.*97G>A | F2 | Thrombophilia | AD | P | MSF | Heterozygous | N | N |
| chr11:46739505 G>A \| c.*97G>A | F2 | Thrombophilia | AD | P | MSF | Heterozygous | N | N |
| chr11:46739505 G>A \| c.*97G>A | F2 | Thrombophilia | AD | P | MSF | Heterozygous | N | N |
| chr11:46739505 G>A \| c.*97G>A | F2 | Thrombophilia | AD | P | MSF | Heterozygous | N | N |
| chr11:46739505 G>A \| c.*97G>A | F2 | Thrombophilia | AD | P | MSF | Heterozygous | N | N |
| chr11:46739505 G>A \| c.*97G>A | F2 | Thrombophilia | AD | P | MSF | Heterozygous | N | N |
| chr11:46739505 G>A \| c.*97G>A | F2 | Thrombophilia | AD | P | MSF | Heterozygous | N | N |
| chr11:46739505 G>A \| c.*97G>A | F2 | Thrombophilia | AD | P | MSF | Heterozygous | N | N |
| chr11:46739505 G>A \| c.*97G>A | F2 | Thrombophilia | AD | P | MSF | Heterozygous | N | N |
| chr11:46739505 G>A \| c.*97G>A | F2 | Thrombophilia | AD | P | MSF | Heterozygous | N | N |
| chr11:46739505 G>A \| c.*97G>A | F2 | Thrombophilia | AD | P | MSF | Heterozygous | N | N |
| chr11:46739505 G>A \| c.*97G>A | F2 | Thrombophilia | AD | P | MSF | Heterozygous | N | N |
| chr11:46739505 G>A \| c.*97G>A | F2 | Thrombophilia | AD | P | MSF | Heterozygous | N | N |
| chr11:46739505 G>A \| c.*97G>A | F2 | Thrombophilia | AD | P | MSF | Heterozygous | N | N |
| chr11:46739505 G>A \| c.*97G>A | F2 | Thrombophilia | AD | P | MSF | Heterozygous | N | N |
| chr11:46739505 G>A \| c.*97G>A | F2 | Thrombophilia | AD | P | MSF | Heterozygous | N | N |
| chr11:46739505 G>A \| c.*97G>A | F2 | Thrombophilia | AD | P | MSF | Heterozygous | N | N |
| chr11:46739505 G>A \| c.*97G>A | F2 | Thrombophilia | AD | P | MSF | Heterozygous | N | N |
| chr11:46739505 G>A \| c.*97G>A | F2 | Thrombophilia | AD | P | MSF | Heterozygous | N | N |
| chr11:46739505 G>A \| c.*97G>A ; chr11:108272785 G>GT \| p.Phe1074fs \| c.3218dupT | F2; ATM | Thrombophilia; Hereditary cancer-predisposing syndrom | AD; AD | P; P | MSF; MSF | Heterozygous; Heterozygous | N; N | N; N |
| chr11:46739505 G>A \| c.*97G>A ; chr11:108335105 T>C \| p.Val2716Ala \| c.8147T>C; chr22:28695868 AG>A \| p.Thr410fs \| c.1229delC; chr11:47337502 G>GA \| p.His831fs \| c.2490dupT | F2; ATM; CHEK2; MYBPC3 | Thrombophilia; Hereditary cancer-predisposing syndrome; Hereditary cancer-predisposing syndrome; Cardiomyopathy | AD; AD; AD; AD | P; P; P; P | MSF; MSF; MSF; MSF | Heterozygous; Heterozygous; Heterozygous; Heterozygous | N; N; N; Y (ECHO) | N; Y; Y; Y |
| chr11:46739505 G>A \| c.*97G>A; chr22:28695219 G>A \| p.Ser471Phe \| c.1412C>T | F2; CHEK2 | Thrombophilia; Hereditary cancer-predisposing syndrome | AD; AD | P; P | MSF; MSF | Heterozygous; Heterozygous | N; N | N; N |

| | | | | | | | | |
|---|---|---|---|---|---|---|---|---|
| chr11:46739505 G>A \| c.*97G>A; chr1:169549811 C>T \| p.Arg534Gln \| c.1601G>A | F2; F5 | Thrombophilia; Thrombophilia | AD; AD | P; P | MSF; MSF | Heterozygous; Heterozygous | N; N | N; N |
| chr11:46739505 G>A \| c.*97G>A; chr1:92263713 C>T \| p.Trp440* \| c.1319G>A | F2; GLMN | Thrombophilia; Glomuvenous malformations | AD; AD | P; LP | MSF; MSF | Heterozygous; Heterozygous | N; N | N; N |
| chr11:46739505 G>A \| c.*97G>A; chr17:37744703 A>C \| p.Val61Gly \| c.182T>G | F2; HNF1B | Thrombophilia; Renal cysts and diabetes syndrome | AD; AD | P; VUS | MSF; VUS-R | Heterozygous; Heterozygous | N; Y (Blood test, Metabolome) | N; N |
| chr11:46739505 G>A \| c.*97G>A; chr8:89947835 T>A \| p.Lys635* \| c.1903A>T | F2; NBN | Thrombophilia; Hereditary cancer-predisposing syndrome | AD; AD | P; P | MSF; MSF | Heterozygous; Heterozygous | N; N | N; Y |
| chr11:46739505 G>A \| c.*97G>A; chr5:96410804 G>GTA \| p.Ser356fs \| c.1063_1064dupTA | F2; PCSK1 | Thrombophilia; Obesity susceptibility | AD; AD | P; LP | MSF; MSF | Heterozygous; Heterozygous | N; N | N; N |
| chr11:46739505 G>A \| c.*97G>A ; chr19:38573304 C>T \| p.Thr4709Met \| c.14126C>T; chr1:241497927 A>ATTT \| p.Lys477dup \| c.1431_1433dupAAA | F2; RYR1; FH | Thrombophilia; Minicore myopathy with external ophthalmoplegia ; Leiomyomatosis and renal cell cancer | AD; AD/AR; AD | P; LP; LP | MSF; MSF; MSF | Heterozygous; Heterozygous; Heterozygous | N; N; N | N; N; N |
| chr1:169549811 C>T \| p.Arg534Gln \| c.1601G>A | F5 | Thrombophilia | AD | P | MSF | Heterozygous | N | N |
| chr1:169549811 C>T \| p.Arg534Gln \| c.1601G>A | F5 | Thrombophilia | AD | P | MSF | Heterozygous | N | N |
| chr1:169549811 C>T \| p.Arg534Gln \| c.1601G>A | F5 | Thrombophilia | AD | P | MSF | Heterozygous | N | N |
| chr1:169549811 C>T \| p.Arg534Gln \| c.1601G>A | F5 | Thrombophilia | AD | P | MSF | Heterozygous | N | N |
| chr1:169549811 C>T \| p.Arg534Gln \| c.1601G>A | F5 | Thrombophilia | AD | P | MSF | Heterozygous | N | N |
| chr1:169549811 C>T \| p.Arg534Gln \| c.1601G>A | F5 | Thrombophilia | AD | P | MSF | Heterozygous | N | N |
| chr1:169549811 C>T \| p.Arg534Gln \| c.1601G>A | F5 | Thrombophilia | AD | P | MSF | Heterozygous | N | N |
| chr1:169549811 C>T \| p.Arg534Gln \| c.1601G>A | F5 | Thrombophilia | AD | P | MSF | Heterozygous | N | N |
| chr1:169549811 C>T \| p.Arg534Gln \| c.1601G>A | F5 | Thrombophilia | AD | P | MSF | Heterozygous | N | N |
| chr1:169549811 C>T \| p.Arg534Gln \| c.1601G>A | F5 | Thrombophilia | AD | P | MSF | Heterozygous | N | N |
| chr1:169549811 C>T \| p.Arg534Gln \| c.1601G>A | F5 | Thrombophilia | AD | P | MSF | Heterozygous | N | N |
| chr1:169549811 C>T \| p.Arg534Gln \| c.1601G>A | F5 | Thrombophilia | AD | P | MSF | Heterozygous | N | N |
| chr1:169549811 C>T \| p.Arg534Gln \| c.1601G>A | F5 | Thrombophilia | AD | P | MSF | Heterozygous | N | N |
| chr1:169549811 C>T \| p.Arg534Gln \| c.1601G>A | F5 | Thrombophilia | AD | P | MSF | Heterozygous | N | N |
| chr1:169549811 C>T \| p.Arg534Gln \| c.1601G>A | F5 | Thrombophilia | AD | P | MSF | Heterozygous | N | N |
| chr1:169549811 C>T \| p.Arg534Gln \| c.1601G>A | F5 | Thrombophilia | AD | P | MSF | Heterozygous | N | N |
| chr1:169549811 C>T \| p.Arg534Gln \| c.1601G>A | F5 | Thrombophilia | AD | P | MSF | Homozygous | N | N |
| chr1:169549811 C>T \| p.Arg534Gln \| c.1601G>A | F5 | Thrombophilia | AD | P | MSF | Heterozygous | N | N |
| chr1:169549811 C>T \| p.Arg534Gln \| c.1601G>A | F5 | Thrombophilia | AD | P | MSF | Heterozygous | N | N |
| chr1:169549811 C>T \| p.Arg534Gln \| c.1601G>A | F5 | Thrombophilia | AD | P | MSF | Heterozygous | N | N |
| chr1:169549811 C>T \| p.Arg534Gln \| c.1601G>A | F5 | Thrombophilia | AD | P | MSF | Heterozygous | N | N |
| chr1:169549811 C>T \| p.Arg534Gln \| c.1601G>A | F5 | Thrombophilia | AD | P | MSF | Heterozygous | N | N |

| | | | | | | | | |
|---|---|---|---|---|---|---|---|---|
| chr1:169549811 C>T \| p.Arg534Gln \| c.1601G>A | F5 | Thrombophilia | AD | P | MSF | Heterozygous | N | N |
| chr1:169549811 C>T \| p.Arg534Gln \| c.1601G>A | F5 | Thrombophilia | AD | P | MSF | Heterozygous | N | N |
| chr1:169549811 C>T \| p.Arg534Gln \| c.1601G>A | F5 | Thrombophilia | AD | P | MSF | Heterozygous | N | N |
| chr1:169549811 C>T \| p.Arg534Gln \| c.1601G>A | F5 | Thrombophilia | AD | P | MSF | Heterozygous | N | N |
| chr1:169549811 C>T \| p.Arg534Gln \| c.1601G>A | F5 | Thrombophilia | AD | P | MSF | Heterozygous | N | N |
| chr1:169549811 C>T \| p.Arg534Gln \| c.1601G>A | F5 | Thrombophilia | AD | P | MSF | Heterozygous | N | N |
| chr1:169549811 C>T \| p.Arg534Gln \| c.1601G>A | F5 | Thrombophilia | AD | P | MSF | Heterozygous | N | N |
| chr1:169549811 C>T \| p.Arg534Gln \| c.1601G>A | F5 | Thrombophilia | AD | P | MSF | Heterozygous | N | N |
| chr1:169549811 C>T \| p.Arg534Gln \| c.1601G>A | F5 | Thrombophilia | AD | P | MSF | Heterozygous | N | N |
| chr1:169549811 C>T \| p.Arg534Gln \| c.1601G>A | F5 | Thrombophilia | AD | P | MSF | Heterozygous | N | N |
| chr1:169549811 C>T \| p.Arg534Gln \| c.1601G>A | F5 | Thrombophilia | AD | P | MSF | Heterozygous | N | N |
| chr1:169549811 C>T \| p.Arg534Gln \| c.1601G>A | F5 | Thrombophilia | AD | P | MSF | Heterozygous | N | N |
| chr1:169549811 C>T \| p.Arg534Gln \| c.1601G>A | F5 | Thrombophilia | AD | P | MSF | Heterozygous | N | N |
| chr1:169549811 C>T \| p.Arg534Gln \| c.1601G>A | F5 | Thrombophilia | AD | P | MSF | Heterozygous | N | N |
| chr1:169549811 C>T \| p.Arg534Gln \| c.1601G>A | F5 | Thrombophilia | AD | P | MSF | Heterozygous | N | N |
| chr1:169549811 C>T \| p.Arg534Gln \| c.1601G>A | F5 | Thrombophilia | AD | P | MSF | Heterozygous | N | N |
| chr1:169549811 C>T \| p.Arg534Gln \| c.1601G>A; chr2:21032391 G>A \| p.Arg439* \| c.1315C>T; chr17:39665813 C>T \| p.Arg70Trp \| c.208C>T | F5; APOB; TCAP | Thrombophilia; Familial hypercholesterolemia; Cardiomyopathy/Muscular dystrophy | AD; AD; AD/AR | P; P; VUS | MSF; MSF; VUS-R | Heterozygous; Heterozygous; Heterozygous | N; N; Y (ECHO, ECG) | N; N; Y |
| chr1:169549811 C>T \| p.Arg534Gln \| c.1601G>A; chr22:28725099 A>G \| p.Ile200Thr \| c.599T>C | F5; CHEK2 | Thrombophilia; Hereditary cancer-predisposing syndrome | AD; AD | P; LP | MSF; MSF | Heterozygous; Heterozygous | N; N | N; Y |
| chr1:169549811 C>T \| p.Arg534Gln \| c.1601G>A; chr22:28725099 A>G \| p.Ile200Thr \| c.599T>C | F5; CHEK2 | Thrombophilia; Hereditary cancer-predisposing syndrome | AD; AD | P; LP | MSF; MSF | Heterozygous; Heterozygous | N; N | N; N |
| chr1:169549811 C>T \| p.Arg534Gln \| c.1601G>A; chr8:19951811 G>A \| p.Ala98Thr \| c.292G>A | F5; LPL | Thrombophilia; Hyperlipidemia | AD; AD | P; LP | MSF; MSF | Heterozygous; Heterozygous | N; Y (Blood test) | N; Y |
| chr1:169549811 C>T \| p.Arg534Gln \| c.1601G>A; chr15:99690400 C>T \| p.Pro277Leu \| c.830C>T | F5; MEF2A | Thrombophilia; Coronary artery disease/myocardial infarction | AD; AD | P; VUS | MSF; VUS-R | Heterozygous; Heterozygous | N; Y (MRI) | N; Y |
| chr1:169549811 C>T \| p.Arg534Gln \| c.1601G>A; chr12:110911176 C>G \| c.403-1G>C | F5; MYL2 | Thrombophilia; Hypertrophic Cardiomyopathy | AD; AD | P; P | MSF; MSF | Heterozygous; Heterozygous | N; N | N; Y |
| chr1:169549811 C>T \| p.Arg534Gln \| c.1601G>A; chr18:31595131 A>G \| p.Glu71Gly \| c.212A>G | F5; TTR | Thrombophilia; Amyloidosis | AD; AD | P; VUS | MSF; VUS-R | Heterozygous; Heterozygous | N; N | N; Y |
| chr1:46405737 G>A \| p.Arg243His \| c.728G>A ; chr1:92271644 T>TA \| p.Leu248fs \| c.743dupT | FAAH; GLMN | Obesity; Glomuvenous malformations | AD; AD | VUS; LP | VUS-R; MSF | Heterozygous; Heterozygous | Y (PHx); N | N; N |
| chr4:83462592 T>TC \| p.Ser370fs \| c.1106dupG | FAM175A | Hereditary cancer-predisposing syndrome | AD | VUS | VUS-R | Heterozygous | Y (PHx, Blood test) | Y |

| | | | | | | | | |
|---|---|---|---|---|---|---|---|---|
| chr1:241497927 A>ATTT \| p.Lys477dup \| c.1431_1433dupAAA | FH | Leiomyomatosis and renal cell cancer | AD | LP | MSF | Heterozygous | N | N |
| chr1:152312600 CACTG>C \| p.Ser761fs \| c.2282_2285delCAGT | FLG | Atopic dermatitis | AD/AR | LP | MSF | Homozygous | N | N |
| chr1:152312600 CACTG>C \| p.Ser761fs \| c.2282_2285delCAGT | FLG | Atopic dermatitis | AD/AR | LP | MSF | Homozygous | N | N |
| chr1:152312600 CACTG>C \| p.Ser761fs \| c.2282_2285delCAGT; chr1:152313655 CGCTGACTGCAGAT>C \| p.Ser407fs \| c.1218_1230delATCTGCAGTCAGC | FLG; FLG | Atopic dermatitis; Atopic dermatitis | AD/AR; AD/AR | LP; LP | MSF; MSF | Heterozygous; Heterozygous | N; N | N; N |
| chr1:152307637 G>A \| p.Gln2417* \| c.7249C>T; chr1:152303356 G>A \| p.Gln3844* \| c.11530C>T; chr12:111803962 G>A \| p.Glu504Lys \| c.1510G>A | FLG; FLG; ALDH2 | Atopic dermatitis; Atopic dermatitis; Alcohol sensitivity | AD/AR; AD/AR; AD | P; LP; P | MSF; MSF; MSF | Heterozygous; Heterozygous; Heterozygous | Y (PHx); Y (PHx); N | N; N; N |
| chr1:171107811 C>T \| p.Pro153Leu \| c.458C>T; chr1:169549811 C>T \| p.Arg534Gln \| c.1601G>A | FMO3; F5 | Trimethylaminuria; Thrombophilia | AR; AD | LP; P | MSF; MSF | Homozygous; Heterozygous | Y (PHx); N | N; N |
| chrX:154532390 G>A \| p.Arg500Cys \| c.1498C>T | G6PD | Glucose 6 phosphate dehydrogenase deficiency | XLD | LP | MSF | Heterozygous | N | N |
| chrX:154532390 G>A \| p.Arg500Cys \| c.1498C>T | G6PD | Glucose 6 phosphate dehydrogenase deficiency | XLD | LP | MSF | Heterozygous | N | N |
| chrX:154536002 C>T \| p.Val68Met \| c.202G>A | G6PD | Glucose 6 phosphate dehydrogenase deficiency | XLD | P | MSF | Hemizygous | Y (PHx) | N |
| chrX:154534419 G>A \| p.Ser188Phe \| c.563C>T | G6PD | Glucose 6 phosphate dehydrogenase deficiency | XLD | LP | MSF | Heterozygous | N | N |
| chrX:154536002 C>T \| p.Val68Met \| c.202G>A | G6PD | Glucose 6 phosphate dehydrogenase deficiency | XLD | P | MSF | Heterozygous | N | N |
| chrX:154532390 G>A \| p.Arg500Cys \| c.1498C>T ; chr11:17407058 G>A \| p.Arg999* \| c.2995C>T | G6PD; ABCC8 | Glucose 6 phosphate dehydrogenase deficiency; Familial hyperinsulinemic hypoglycemia | XLD; AD/AR | LP; LP | MSF; MSF | Hemizygous; Heterozygous | N; Y (Blood test, Metabolome) | N; Y |
| chr2:17761537 G>A \| p.Trp101* \| c.303G>A | GEN1 | Cancer-predisposing syndrome | AD | VUS | VUS-R | Heterozygous | Y (PHx) | Y |
| chr2:17760056 CACAG>C \| p.Thr40fs \| c.118_121delACAG | GEN1 | Cancer-predisposing syndrome | AD | VUS | VUS-R | Heterozygous | N | Y |
| chr13:20189481 A>G \| p.Met34Thr \| c.101T>C; chr13:20189546 AC>A \| p.Gly12fs \| c.35delG | GJB2; GJB2 | Deafness; Deafness | AR; AR | P; P | MSF; MSF | Heterozygous; Heterozygous | N; N | N; N |
| chr13:20189346 AG>A \| p.Leu79fs \| c.235delC; chr13:20189473 C>T \| p.Val37Ile \| c.109G>A; chr12:111803962 G>A \| p.Glu504Lys \| c.1510G>A | GJB2; GJB2; ALDH2 | Deafness; Deafness; Alcohol sensitivity | AR; AR; AD | P; LP; P | MSF; MSF; MSF | Heterozygous; Heterozygous; Homozygous | N; N; N | N; N; N |
| chr3:32159096 C>T \| p.Ala280Val \| c.839C>T | GPD1L | Brugada syndrome | AD | LP | MSF | Heterozygous | N | N |
| chr3:32138608 G>A \| p.Glu83Lys \| c.247G>A | GPD1L | Brugada syndrome | AD | VUS | VUS-R | Heterozygous | Y (iRhythm) | N |
| chr17:44351750 C>A \| p.Cys378* \| c.1134C>A | GRN | Frontotemporal lobar degeneration with ubiquitin-positive inclusions | AD | LP | MSF | Heterozygous | N | N |
| chr11:5227002 T>A \| p.Glu7Val \| c.20A>T | HBB | Beta thalassemia | AD/AR | P | MSF | Heterozygous | Y (Blood test) | N |
| chr11:5225678 C>G \| c.364G>C | HBB | Beta thalassemia | AD/AR | P | MSF | Heterozygous | Y (Blood test) | N |

| | | | | | | | | |
|---|---|---|---|---|---|---|---|---|
| chr11:5226985 TA>T \| p.Thr13fs \| c.36delT | HBB | Beta thalassemia | AD/AR | P | MSF | Heterozygous | Y (PHx) | N |
| chr11:5226995 CTT>C \| p.Lys9fs \| c.25_26delAA | HBB | Beta thalassemia | AD/AR | LP | MSF | Heterozygous | Y (Blood test) | N |
| chr11:5227003 CAG>C \| p.Pro6fs \| c.17_18delCT ; chr17:37744860 G>A \| p.Gln9* \| c.25C>T | HBB; HNF1B | Beta thalassemia; Renal cysts and diabetes syndrome | AD/AR; AD | P; LP | MSF; MSF | Heterozygous; Heterozygous | Y (PHx); Y (MRI) | N; N |
| chr6:26092913 G>A \| p.Cys282Tyr \| c.845G>A | HFE | Hemochromatosis | AR | P | MSF | Homozygous | Y (MRI) | N |
| chr6:26092913 G>A \| p.Cys282Tyr \| c.845G>A | HFE | Hemochromatosis | AR | P | MSF | Homozygous | Y (MRI, Blood test) | N |
| chr12:120994313 G>GC \| p.Pro289fs \| c.863_864insC | HNF1A | Maturity-onset diabetes of the young, type 3 | AD | LP | MSF | Heterozygous | Y (Blood test, Metabolome) | Y |
| chr17:48728343 C>T \| p.Gly84Glu \| c.251G>A | HOXB13 | Cancer-predisposing syndrome | AD | LP | MSF | Heterozygous | N | N |
| chr17:48728343 C>T \| p.Gly84Glu \| c.251G>A | HOXB13 | Cancer-predisposing syndrome | AD | LP | MSF | Heterozygous | N | N |
| chr17:48728343 C>T \| p.Gly84Glu \| c.251G>A | HOXB13 | Cancer-predisposing syndrome | AD | LP | MSF | Heterozygous | N | N |
| chr17:48728343 C>T \| p.Gly84Glu \| c.251G>A | HOXB13 | Cancer-predisposing syndrome | AD | LP | MSF | Heterozygous | N | N |
| chr17:48728343 C>T \| p.Gly84Glu \| c.251G>A | HOXB13 | Cancer-predisposing syndrome | AD | LP | MSF | Heterozygous | N | N |
| chr17:48728343 C>T \| p.Gly84Glu \| c.251G>A; chr4:89828156 A>C \| p.His50Gln \| c.150T>G | HOXB13; SNCA | Cancer-predisposing syndrome; Dementia/Parkinson disease | AD; AD | LP ; VUS | MSF; VUS-R | Heterozygous; Heterozygous | N; N | N; Y |
| chr19:7184486 G>T \| p.Cys268* \| c.804C>A | INSR | Insulin resistance/Leprechaunism | AD | LP | MSF | Heterozygous | Y (Metabolome) | N |
| chr17:47300488 G>T \| p.Glu642* \| c.1924G>T ; chr17:47253911 TG>T \| p.Ala19fs \| c.55delG; chr17:16948947 T>C \| p.Tyr79Cys \| c.236A>G | ITGB3; ITGB3; TNFRSF13B | Glanzmann's thrombasthenia ; Glanzmann's thrombasthenia ; Common variable immune deficiency | AR; AR; AD/AR | P; LP; VUS | MSF; MSF; VUS-R | Heterozygous; Heterozygous; Heterozygous | N; N; Y (PHx) | N; N; N |
| chr21:34370639 T>C \| p.Met54Thr \| c.161T>C | KCNE2 | Long QT syndrome | AD | LP | MSF | Heterozygous | N | N |
| chr11:2583439 C>T \| p.Thr309Ile \| c.926C>T | KCNQ1 | Long QT syndrome | AD | LP | MSF | Heterozygous | Y (ECG) | N |
| chr11:2572871 G>A \| p.Gly269Asp \| c.806G>A | KCNQ1 | Long QT syndrome | AD | P | MSF | Heterozygous | N | Y |
| chr2:8802974 G>A \| p.Gln253* \| c.757C>T | KIDINS220 | Spastic paraplegia, intellectual disability, nystagmus, and obesity | AD | LP | MSF | Heterozygous | N | N |
| chr19:11113557 G>A \| p.Gly461Ser \| c.1381G>A | LDLR | Familial hypercholesterolemia | AD | VUS | VUS-R | Heterozygous | Y (Blood test) | N |
| chr19:11089428 T>C \| c.-121T>C | LDLR | Familial hypercholesterolemia | AD | LP | MSF | Heterozygous | Y (Blood test) | N |
| chr19:11113590 G>T \| p.Asp472Tyr \| c.1414G>T | LDLR | Familial hypercholesterolemia | AD | LP | MSF | Heterozygous | Y (Blood test) | Y |
| chr19:11113557 G>A \| p.Gly461Ser \| c.1381G>A; chr3:15645186 G>C \| p.Asp446His \| c.1336G>C | LDLR; BTD | Familial hypercholesterolemia ; Biotinidase deficiency | AD; AR | VUS; LP | VUS-R; MSF | Heterozygous; Homozygous | Y (Blood test); N | Y; N |
| chr19:11110658 A>G \| p.Asn316Ser \| c.947A>G; chr11:5226943 C>T \| c.79G>A | LDLR; HBB | Familial hypercholesterolemia; Beta thalassemia | AD; AD/AR | LP; P | MSF; MSF | Heterozygous; Heterozygous | Y (Blood test); Y (Blood test) | Y; N |
| chr1:156130658 G>A \| p.Arg133Gln \| c.398G>A | LMNA | Cardiomyopathy | AD/AR | LP | MSF | Heterozygous | Y (ECHO, ECG) | Y |
| chr8:19948181 AAGAAG>A \| p.Arg32fs \| c.94_98delAGAGA | LPL | Hyperlipidemia | AD | LP | MSF | Heterozygous | Y (PHx) | Y |
| chr8:19951805 G>C \| p.Val96Leu \| c.286G>C | LPL | Hyperlipidemia | AD | VUS | VUS-R | Heterozygous | Y (Blood test) | Y |

| | | | | | | | | |
|---|---|---|---|---|---|---|---|---|
| chr11:68413958 C>T \| p.Arg925Cys \| c.2773C>T; chr1:149784224 C>T \| p.Arg92* \| c.274C>T | LRP5; FCGR1A | Osteoporosis ; IGG receptor I, phagocytic, familial deficiency of | AD; AD | VUS; LP | VUS-R; MSF | Heterozygous; Heterozygous | Y (PHx); N | N; N |
| chr12:40340400 G>A \| p.Gly2019Ser \| c.6055G>A | LRRK2 | Parkinson disease | AD | P | MSF | Heterozygous | N | N |
| chr12:40235634 T>C \| p.Leu119Pro \| c.356T>C | LRRK2 | Parkinson disease | AD | VUS | VUS-R | Heterozygous | N | Y |
| chr22:20983063 C>CA \| p.Ile80fs \| c.238dupA | LZTR1 | Schwannomatosis | AD | LP | MSF | Heterozygous | N | N |
| chr22:20996723 C>A \| p.Tyr749* \| c.2247C>A | LZTR1 | Schwannomatosis | AD | LP | MSF | Heterozygous | N | N |
| chr22:20991813 CCAGCTCCG>C \| p.Ser327fs \| c.978_985delCAGCTCC | LZTR1 | Schwannomatosis | AD | LP | MSF | Heterozygous | N | N |
| chr22:20992304 C>T \| p.Arg362* \| c.1084C>T; chr15:99690400 C>T \| p.Pro277Leu \| c.830C>T | LZTR1; MEF2A | Schwannomatosis; Coronary artery disease/myocardial infarction | AD; AD | LP; VUS | MSF; VUS-R | Heterozygous; Heterozygous | N; N | N; Y |
| chr2:19997171 AG>A \| p.Pro419fs \| c.1256delC | MATN3 | Osteoarthritis susceptibility 2 | AD | VUS | VUS-R | Heterozygous | Y (PHx) | Y |
| chr18:60371842 T>C \| p.Ile170Val \| c.508A>G | MC4R | Obesity | AD | LP | MSF | Heterozygous | Y (PHx) | N |
| chr11:60093404 G>GAGGA \| p.Ser130fs \| c.386_389dupGAAG | MS4A2 | Atopy | AD | VUS | VUS-R | Heterozygous | Y (PHx) | N |
| chr2:47783243 C>T \| p.Gln4* \| c.10C>T | MSH6 | Lynch syndrome | AD | LP | MSF | Heterozygous | N | N |
| chr11:47342734 C>T \| p.Gly490Arg \| c.1468G>A | MYBPC3 | Cardiomyopathy | AD | VUS | VUS-R | Heterozygous | Y (PHx, ECHO) | Y |
| chr11:47332274 AGAGAGGGAGGGAAGCCATCCAGGCT>A \| c.*3125_*3149delGAGAGGGAGGGAAGCCATCCAGGCT | MYBPC3 | Cardiomyopathy | AD | LP | MSF | Heterozygous | Y (iRhythm) | Y |
| chr11:47332658 C>T \| p.Glu1179Lys \| c.3535G>A | MYBPC3 | Cardiomyopathy | AD | VUS | VUS-R | Heterozygous | Y (ECHO) | Y |
| chr11:47342698 G>A \| p.Arg502Trp \| c.1504C>T | MYBPC3 | Cardiomyopathy | AD | P | MSF | Heterozygous | Y (ECHO) | N |
| chr11:47337722 G>A \| p.Pro794Leu \| c.2381C>T | MYBPC3 | Cardiomyopathy | AD | VUS | VUS-R | Heterozygous | Y (ECHO) | N |
| chr7:155803420 C>T \| p.Gly290Asp \| c.869G>A | MYBPC3 | Cardiomyopathy | AD | VUS | VUS-R | Heterozygous | N | Y |
| chr11:47347891 C>T \| p.Gly263Arg \| c.787G>A | MYBPC3 | Cardiomyopathy | AD | VUS | VUS-R | Heterozygous | N | Y |
| chr11:47346297 C>T \| p.Glu334Lys \| c.1000G>A; chr22:28695858 G>A \| p.His414Tyr \| c.1240C>T | MYBPC3; CHEK2 | Cardiomyopathy; Hereditary cancer-predisposing syndrome | AD; AD | VUS; VUS | VUS-R; VUS-R | Heterozygous; Heterozygous | Y (ECHO); N | Y; Y |
| chr14:23424875 C>T \| p.Arg858His \| c.2573G>A | MYH7 | Cardiomyopathy | AD | VUS | MSF | Heterozygous | Y (ECHO) | Y |
| chr14:23431468 C>T \| p.Arg249Gln \| c.746G>A | MYH7 | Cardiomyopathy | AD | P | MSF | Heterozygous | Y (ECHO) | Y |
| chr14:23413859 C>T \| p.Arg1897His \| c.5690G>A | MYH7 | Cardiomyopathy | AD | VUS | VUS-R | Heterozygous | Y (ECHO, ECG) | Y |
| chr14:23433704 C>G \| p.Gly10Ala \| c.29G>C; chr6:26092913 G>A \| p.Cys282Tyr \| c.845G>A | MYH7; HFE | Cardiomyopathy; Hemochromatosis | AD; AR | VUS; P | VUS-R; MSF | Heterozygous; Homozygous | Y (ECHO, ECG); Y (PHx) | Y; Y |
| chr12:110919160 C>T \| p.Ala13Thr \| c.37G>A | MYL2 | Hypertrophic Cardiomyopathy | AD | VUS | VUS-R | Heterozygous | Y (ECHO) | Y |
| chr12:110911176 C>G \| c.403-1G>C | MYL2 | Hypertrophic Cardiomyopathy | AD | P | MSF | Heterozygous | N | Y |

| | | | | | | | | |
|---|---|---|---|---|---|---|---|---|
| chr12:110913097 T>G \| p.Glu134Ala \| c.401A>C; chr5:13919318 GC>G \| p.Ala278fs \| c.832delG; chr5:13791994 ATTTGGTTC>A \| p.Glu2814fs \| c.8440_8447delGAACCAAA | MYL2; DNAH5; DNAH5 | Hypertrophic Cardiomyopathy; Ciliary dyskinesia; Ciliary dyskinesia | AD; AR; AR | VUS; LP; LP | VUS-R; MSF; MSF | Heterozygous; Heterozygous; Heterozygous | Y (ECHO); N; N | N; N; N |
| chr3:123707995 C>A \| p.Asp717Tyr \| c.2149G>T | MYLK | Thoracic aortic aneurysm and aortic dissection | AD | VUS | VUS-R | Heterozygous | Y (ECHO) | N |
| chr3:123708870 C>A \| p.Trp656Cys \| c.1968G>T | MYLK | Thoracic aortic aneurysm and aortic dissection | AD | VUS | VUS-R | Heterozygous | Y (ECHO) | N |
| chr17:18119933 TC>T \| p.Tyr380fs \| c.1137delC; chr17:18121804 AC>A \| p.Lys1003fs \| c.3006delC | MYO15A; MYO15A | Deafness; Deafness | AR; AR | LP; LP | MSF; MSF | Heterozygous; Heterozygous | N; N | N; N |
| chr8:89955524 C>CT \| p.Val386fs \| c.1155dupA | NBN | Hereditary cancer-predisposing syndrome | AD | LP | MSF | Heterozygous | N | Y |
| chr8:89971213 ATTTGT>A \| p.Lys219fs \| c.657_661delACAAA | NBN | Hereditary cancer-predisposing syndrome | AD | P | MSF | Heterozygous | N | Y |
| chr8:89982766 G>A \| p.Arg43* \| c.127C>T | NBN | Hereditary cancer-predisposing syndrome | AD | P | MSF | Heterozygous | N | Y |
| chr2:181678872 C>T \| c.-11-1G>A | NEUROD1 | Maturity-onset diabetes of the young 6 | AD | LP | MSF | Heterozygous | Y (Metabolome) | Y |
| chr17:31261733 C>T \| p.Arg1534* \| c.4600C>T | NF1 | Neurofibromatosis | AD | P | MSF | Heterozygous | Y (MRI) | N |
| chr1:9982570 C>T \| p.Arg237Cys \| c.709C>T | NMNAT1 | Leber congenital amaurosis 9 | AR | LP | MSF | Heterozygous | N | N |
| chr1:9982577 T>C \| p.Leu239Ser \| c.716T>C | NMNAT1 | Leber congenital amaurosis 9 | AR | LP | MSF | Heterozygous | N | N |
| chr7:144404628 G>T \| p.Tyr46* \| c.138C>A | NOBOX | Premature ovarian failure/Familial cancer of breast | AD | LP | MSF | Heterozygous | N | N |
| chr7:44517253 G>A \| p.Arg1108Trp \| c.3322C>T | NPC1L1 | Low density lipoprotein cholesterol level QTL 7 | AD | LP | MSF | Heterozygous | Y (Blood test) | N |
| chr5:143300673 A>AT \| p.Ile521fs \| c.1561dupA | NR3C1 | Glucocorticoid resistance | AD | LP | MSF | Heterozygous | Y (Blood test) | N |
| chr11:117171692 C>T \| c.*3993C>T | PAFAH1B2 | Increased HDL cholesterol levels | AD | VUS | VUS-R | Heterozygous | Y (Blood test) | N |
| chr16:23636038 T>C \| p.Arg170Gly \| c.508A>G | PALB2 | Familial cancer of breast, Hereditary cancer-predisposing syndrome | AD | VUS | VUS-R | Heterozygous | Y (PHx) | N |
| chr5:96408289 G>A \| p.Thr377Met \| c.1130C>T | PCSK1 | Obesity susceptibility | AD | VUS | VUS-R | Heterozygous | Y (PHx) | N |
| chr1:55052398 G>A \| p.Arg215His \| c.644G>A | PCSK9 | Familial hypercholesterolemia | AD | P | MSF | Heterozygous | Y (PHx) | N |
| chr1:55058543 C>G \| c.1399C>G | PCSK9 | Familial hypercholesterolemia | AD | VUS | VUS-R | Heterozygous | Y (PHx) | Y |
| chr1:55039847 G>A \| p.Val4Ile \| c.10G>A; chr12:111803962 G>A \| p.Glu504Lys \| c.1510G>A | PCSK9; ALDH2 | Familial hypercholesterolemia; Alcohol sensitivity | AD; AD | P; P | MSF; MSF | Heterozygous; Heterozygous | Y (MRI, Blood test); N | N; N |
| chr16:2110286 A>T \| p.Tyr1627* \| c.4881T>A | PKD1 | Polycystic kidney disease | AD | LP | MSF | Heterozygous | Y (MRI, Blood test, Metabolome) | Y |
| chr12:32878941 AG>A \| p.Pro105fs \| c.314delC | PKP2 | Arrhythmogenic right ventricular dysplasia 9 | AD | P | MSF | Heterozygous | Y (ECHO, ECG) | Y |
| chr2:189863774 C>T \| p.Arg630* \| c.1888C>T | PMS1 | Lynch syndrome | AD | VUS | VUS-R | Heterozygous | Y (PHx) | Y |
| chr7:5987534 C>A \| p.Glu411* \| c.1231G>T | PMS2 | Lynch syndrome | AD | LP | MSF | Heterozygous | N | Y |
| chr7:113879105 CCT>C \| p.Gln662fs \| c.1985_1986delAG | PPP1R3A | Diabetes mellitus, type II, Insulin resistance, severe, | AD | LP | MSF | Heterozygous | N | N |
| chr7:113879105 CCT>C \| p.Gln662fs \| c.1985_1986delAG | PPP1R3A | Diabetes mellitus, type II, Insulin resistance, severe | AD | LP | MSF | Heterozygous | Y (PHx) | N |

| | | | | | | | | |
|---|---|---|---|---|---|---|---|---|
| chr2:127426208 G>A \| p.Arg254Gln \| c.761G>A | PROC | Protein C deficiency | AD | LP | MSF | Heterozygous | N | N |
| chr2:127426114 C>T \| p.Arg223Trp \| c.667C>T ; chr12:111803962 G>A \| p.Glu504Lys \| c.1510G>A | PROC; ALDH2 | Protein C deficiency; Alcohol sensitivity | AD; AD | LP; P | MSF; MSF | Heterozygous; Heterozygous | N; N | N; N |
| chr3:71781525 AT>A \| p.Ile55fs \| c.163delA | PROK2 | Hypogonadotropic hypogonadism 4 with or without anosmia | AD | LP | MSF | Heterozygous | N | N |
| chr19:54123818 CTCCAAGCACCGCA>C \| p.Lys201fs \| c.600_612delCAAGCACCGCATC | PRPF31 | Retinitis pigmentosa | AD | LP | MSF | Heterozygous | N | N |
| chr5:132595780 G>A \| p.Arg726His \| c.2177G>A | RAD50 | Hereditary cancer-predisposing syndrome | AD | VUS | VUS-R | Heterozygous | Y (PHx) | Y |
| chr5:132575885 AAGAC>A \| p.Thr109fs \| c.326_329delCAGA | RAD50 | Hereditary cancer-predisposing syndrome | AD | P | MSF | Heterozygous | N | Y |
| chr17:58696857 AG>A \| c.571+1delG | RAD51C | Breast-ovarian cancer, familial, susceptibility to, 3 | AD | LP | MSF | Heterozygous | N | N |
| chr11:47448079 G>T \| p.Asn88Lys \| c.264C>A | RAPSN | Myasthenic syndrome | AR | P | MSF | Homozygous | Y (PHx) | Y |
| chr13:48459708 C>T \| c.1981C>T | RB1 | Retinoblastoma | AD | P | MSF | Heterozygous | N | N |
| chr10:110781125 AC>A \| p.Ser175fs \| c.522delC | RBM20 | Cardiomyopathy | AD | LP | MSF | Heterozygous | Y (ECHO) | N |
| chr12:21483433 G>A \| p.Arg215* \| c.643C>T | RECQL | Hereditary cancer-predisposing syndrome | AD | LP | MSF | Heterozygous | N | Y |
| chr1:237454494 C>G \| p.Pro466Ala \| c.1396C>G | RYR2 | Ventricular tachycardia, polymorphic | AD | VUS | VUS-R | Heterozygous | Y (ECHO) | Y |
| chr2:166012122 A>G \| p.Phe1289Ser \| c.3866T>C; chr4:186280258 T>C \| p.Phe301Leu \| c.901T>C | SCN1A; F11 | Epilepsy; Factor XI deficiency | AD; AD/AR | VUS; P | VUS-R; MSF | Heterozygous; Heterozygous | Y (PHx); N | Y; N |
| chr19:35033545 G>A \| p.Arg85His \| c.254G>A | SCN1B | Atrial fibrillation, epilepsy | AD | LP | MSF | Heterozygous | Y (ECG) | N |
| chr14:94380925 T>A \| p.Glu288Val \| c.863A>T; chr14:94378610 C>T \| p.Glu366Lys \| c.1096G>A | SERPINA1; SERPINA1 | Alpha-1 antitrypsin deficiency; Alpha-1 antitrypsin deficiency | AR; AR | LP; P | MSF; MSF | Heterozygous; Heterozygous | N; N | N; N |
| chr7:94656078 C>T \| p.Trp7* \| c.21G>A | SGCE | Myoclonus dystonia | AD | LP | MSF | Heterozygous | N | N |
| chr5:132369824 G>A \| c.-149G>A | SLC22A5 | Carnitine deficiency | AR | VUS | VUS-R | Heterozygous | N | N |
| chr15:66781472 C>G \| p.Tyr476* \| c.1428C>G | SMAD6 | Aortic valve disease | AD | VUS | MSF | Heterozygous | Y (PHx) | N |
| chr13:75362451 AG>A \| p.Ser219fs \| c.654delC | TBC1D4 | Diabetes mellitus, noninsulin-dependent | AD | VUS | VUS-R | Heterozygous | Y (Metabolome, Blood test, PHx) | Y |
| chr20:2417241 AAG>A \| p.Arg451fs \| c.1351_1352delAG | TGM6 | Spinocerebellar ataxia | AD | VUS | VUS-R | Heterozygous | Y (PHx) | N |
| chr1:201359245 G>A \| p.Arg288Cys \| c.862C>T | TNNT2 | Cardiomyopathy | AD | LP | MSF | Heterozygous | Y (ECHO) | N |
| chr17:7673776 G>A \| c.844C>T | TP53 | Osteosarcoma, Li Fraumeni-like syndrome | AD | LP | MSF | Heterozygous | Y (PHx) | Y |
| chr2:178588009 G>GT \|p.Thr18565fs \| c.55693dupA | TTN | Cardiomyopathy | AD | LP | MSF | Heterozygous | N | N |
| chr2:178572286 G>A \| p.Arg24616* \| c.73846C>T | TTN | Cardiomyopathy | AD | LP | MSF | Heterozygous | N | N |
| chr18:31598655 G>A \| p.Val180Ile \| c.538G>A | TTR | Amyloidosis | AD | P | MSF | Heterozygous | Y (PHx) | N |
| chr4:6301467 C>T \| p.Arg558Cys \| c.1672C>T | WFS1 | Wolfram syndrome | AR | VUS | VUS-R | Heterozygous | Y (Blood test, Metabolome) | Y |

Abbreviations: MOI, mode of inheritance; ACMG, American College of Medical Genomics and Genetics; AD, autosomal dominant; AR, autosomal recessive; XLD, x-linked dominant; P, pathogenic; LP, likely pathogenic; VUS, variant of unknown significance; MSF, medically significant genetic findings; VUS-R, reportable variant of unknown significance; Y, Yes; N, No.

**Supplemental Table S4A: Extreme Metabolites (±6 S.D.) and Associated Genetic Variants.**

| Metabolite | Gene | Variant (Number of cases) |
|---|---|---|
| Arabitol-xylitol (↑) | *AKR1C3* | c.584C>T (1) |
| Taurodeoxycholate (↓) | *AKR1C4* | c.434C>G(;)931C>G (1[#]) |
| Adenosine 5'-monophosphate (AMP) (↓) | *AMPD1* | c.133C>T (5, three heterozygotes, two homozygotes)<br>c.1820G>A (1)<br>c.323G>A (1)<br>c.959A>T (1) |
| Alpha-hydroxy isovalerate (↓) | *BCAT1* | c.212C>T (1) |
| Aamma-glutamyl-epsilon-lysine (↓), 5-oxoproline (↓) | *CHAC1* | c.-53A>C (1)<br>c.-806C>G (1) |
| 1-palmitoyl-2-adrenoyl-GPC (16:0/22:4) (↓) | *CHKA* | c.-1529delA (1) |
| 1-2-dilinoleoyl-GPC (18:2/18:2) (↓), 1-adrenoyl-GPC (22:4) (↓) | *CHKB* | c.-19G>T(;)27G>A (1[#]) |
| Theobromine (↓) | *CYP1A2* | c.1459G>A (1) |
| Cortisol (↓) | *CYP21A2* | c.1179C>G (1) |
| Dimethylglycine (↑) | *DMGDH* | c.1300A>T(;)1747G>T* (1[#]) |
| Gamma-glutamylvaline (↓) | *GGCT* | c.141+137G>A (1) |
| Glutamate (↓) | *GLS* | c.-557C>T (1) |
| Glutamate (↓) | *GLS2* | c.-262C>A (2) |
| Glutamate (↓) | *GLUD2* | c.-922T>A(;)1492T>G (1[#]) |
| Serotonin (↓) | *MAOA* | c.-1227_-1226ins30 (1)<br>c.306+2836A>G (1 hemizygote) |
| 5-oxoproline (↓) | *OPLAH* | c.1502G>A (1) |
| 5-oxoproline (↓) | *OPLAH* | c.-728G>C (1) |
| 1-palmitoyl-2-adrenoyl-GPC (16:0/22:4) (↓) | *PC* | c.-1+5138G>A (1) |
| 1-1-enyl-oleoyl-GPE (P-18:1) (↓) | *PLA2G2A* | c.332A>G (1) |
| 1-1-enyl-oleoyl-GPE (P-18:1) (↓) | *PLA2G3* | c.143T>C (1) |
| 1-1-enyl-oleoyl-GPE (P-18:1) (↓) | *PLA2G4D* | c.46G>C (1) |
| Succinate (↓) | *SDHC* | c.-143G>A (1) |
| Dehydroisoandrosterone sulfate (DHEA-S) (↓) | *SULT2B1* | c.98G>A (1) |
| Alpha-tocopherol (↓) | *TTPA* | c.13C>T[$] (1)<br>c.513_514insTT[$] (1) |

Variants are in heterozygous state unless noted.

*: Reported as medically significant finding (MSF) or reportable variant of uncertain significance (VUS-R).

#: Potential compound heterozygote, but phase of variants is undetermined.

$: Reported as likely pathogenic recessive variant in the carrier.

### Supplemental Table S4B: Pathogenic/Likely Pathogenic Variants Affecting Serum Metabolite Level.

| Gene | Variant (Number of cases) | Elevated/decreased metabolites (95% CI) |
|---|---|---|
| *ACADM* | c.1084A>G (2) | Medium chain fatty acids (↑) |
| *ACADS* | c.319C>T (1) | Ethylmalonate (↑) |
| *ACADSB* | c.621G>A (1) | 2-methylbutyrylcarnitine-C5- (↑) |
| *ACY1* | c.575dupG (1) | N-acetylmethionine, N-acetylvaline, N-acetylalanine, N-acetylglutamate, N-acetylglycine and N-acetylserine (↑) |
| *APOB* | c.13480_13482delCAG (1) | Cholesterol (↑) |
| *APOB* | c.8912A>C (1) | Cholesterol (↑) |
| *ASS1* | c.1030C>T (1) | Citrulline (↑) |
| *CPT2* | c.338C>T (1) | Long chain fatty acids (↑) |
| *CTH* | c.200C>T (2) | Cystathionine (↑) |
| *DMGDH* | c.1300A>T(;)1747G>T* (1[#]) | Dimethylglycine (↑) |
| *HMGCL* | c.853delC (1) | 3-hydroxy-3-methylglutarate (↑) |
| *HOGA1* | c.944_946delAGG (1) | Oxalate (↑) |
| *LDLR* | c.*2117C>G (1) | Cholesterol (↑) |
| *MVK* | c.1129G>A (1) | 3-hydroxy-3-methylglutarate (↑) |
| *NR3C1* | c.1561dupA (1) | Cortisol (↑) |
| *PAH* | c.1139C>T (2) | Phenylalanine (↑) |
| | c.1208C>T (1) | |
| | c.1222C>T (2) | |
| | c.194T>C (1) | |
| | c.611A>G (1) | |
| | c.782G>A (1) | |
| | c.814G>T (1) | |
| | c.829T>G (1) | |
| *PRODH* | c.1322T>C (1) | Proline (↑) |
| *SUGCT* | c.1006C>T (1) | Glutarate (↑) |
| *TTPA* | c.13C>T (3) | Alpha-tocopherol (↓) |
| | c.513_514insTT (1) | |

Variants are in heterozygous status unless noted.

*: Reported as medically significant finding (MSF) or reportable variant of uncertain significance (VUS-R).

#: Potential compound heterozygote, but phase of variants is undetermined.

### Supplemental Table S5: Genome-wide Gene-based Collapsing Analysis.

| Gene | Known condition | P value (all participants) | P value (European ancestry) | Direction of effect | Metabolite | Carrier frequency (cohort in this study combined with TwinsUK cohort) |
|---|---|---|---|---|---|---|
| *PAH* | Phenylketonuria | 2.0E-08 | 3.1E-12 | ↑ | phenylalanine | 0.88% |
| *ETFDH* | Glutaric acidemia | 3.4E-08 | 4.7E-08 | ↑ | X - 12459 | 0.17% |
| *PAH* | Phenylketonuria | 7.2E-07 | 1.9E-08 | ↑ | gamma-glutamylphenylalanine | 0.88% |
| *ETFDH* | Glutaric acidemia | 1.6E-06 | 1.3E-07 | ↑ | octanoylcarnitine | 0.17% |
| *ETFDH* | Glutaric acidemia | 2.3E-06 | 2.3E-07 | ↑ | decanoylcarnitine | 0.17% |
| *DMGDH* | Dimethylglycine dehydrogenase deficiency | 1.4E-43 | 4.5E-31 | ↑ | dimethylglycine | 1.41% |
| *ABHD14A-ACY1* | Aminoacylase 1 deficiency | 7.7E-23 | 1.1E-17 | ↑ | N-acetylalanine | 0.68% |
| *ABHD14A-ACY1* | Aminoacylase 1 deficiency | 2.2E-22 | 3.8E-18 | ↑ | N-acetylmethionine | 0.68% |
| *ABHD14A-ACY1* | Aminoacylase 1 deficiency | 3.6E-18 | 1.1E-13 | ↑ | N-formylmethionine | 0.68% |
| *SLC5A10* | None | 1.7E-17 | 4.9E-20 | ↓ | 1,5-anhydroglucitol (1,5-AG) | 1.03% |
| *ABHD14A-ACY1* | Aminoacylase 1 deficiency | 7.8E-16 | 4.6E-20 | ↑ | N-acetylserine | 0.68% |
| *SLC6A19* | Hyperglycinuria | 1.0E-14 | 2.9E-12 | ↓ | methionine sulfone | 0.95% |
| *HAL* | Histidinemia | 1.0E-12 | 1.5E-09 | ↑ | histidine | 1.62% |
| *APOC3* | Apolipoprotein C-III deficiency | 2.1E-11 | 3.1E-11 | ↓ | 1-palmitoyl-2-arachidonoyl-GPE (16:0/20:4) | 0.29% |
| *HAO2* | None | 6.1E-11 | 1.2E-11 | ↑ | alpha-hydroxyisovalerate | 0.67% |
| *APOC3* | Apolipoprotein C-III deficiency | 9.2E-11 | 1.4E-10 | ↓ | 1-stearoyl-2-arachidonoyl-GPE (18:0/20:4) | 0.29% |
| *APOC3* | Apolipoprotein C-III deficiency | 1.8E-10 | 2.1E-10 | ↓ | 1-stearoyl-2-linoleoyl-GPE (18:0/18:2) | 0.29% |
| *APOC3* | Apolipoprotein C-III deficiency | 2.1E-10 | 3.0E-10 | ↓ | 1-palmitoyl-2-docosahexaenoyl-GPE (16:0/22:6) | 0.29% |
| *HYKK* | None | 4.7E-10 | 1.7E-16 | ↑ | 5-hydroxylysine | 0.67% |
| *ALPL* | Hypophosphatasia | 8.8E-10 | 3.5E-09 | ↑ | choline phosphate | 0.49% |
| *APOC3* | Apolipoprotein C-III deficiency | 1.1E-09 | 1.2E-09 | ↓ | 1-stearoyl-2-docosahexaenoyl-GPE (18:0/22:6) | 0.29% |
| *NIT2* | None | 1.2E-09 | 1.1E-09 | ↑ | alpha-ketoglutaramate | 0.34% |
| *PTER* | None | 1.8E-09 | 3.9E-13 | ↑ | N-acetyl-beta-alanine | 0.51% |

| | | | | | | |
|---|---|---|---|---|---|---|
| *HAO2* | None | 2.4E-09 | 5.6E-10 | ↑ | 2-hydroxy-3-methylvalerate | 0.67% |

### Supplemental Table S6: Identification of Short Tandem Repeats in the Cohort.

| Disease | Gene | Motif | Inheritance | Risk cutoff | Number of At-Risk Individuals |
|---|---|---|---|---|---|
| Susceptibility to prostate cancer due to AR expression (MIM: 176807) | *AR* (MIM: 313700) | CAG | XLR | ≤17 (pre-risk) | 29 |
| Susceptibility to prostate cancer due to AR expression (MIM: 176807) | *AR* (MIM: 313700) | CAG | XLR | ≤8 (risk) | 3 |
| Central hypoventilation syndrome (MIM: 209880, CCHS) | *PHOX2B* (MIM: 603851) | GCN | AD | ≥24 | 1 |
| Myotonic dystrophy 2 (MIM: 602668, DM2) | *ZNF9* (MIM: 116955) | CCTG | AD | ≥75 | 2 |
| Spinal and bulbar muscular atrophy of Kennedy (MIM: 313200, SBMA) | *AR* (MIN: 313700) | CAG | XLR | ≥36 | 1 |
| Spinocerebellar ataxia 17 (MIM: 607136, SCA17) | *TBP* (MIM: 600075) | CAG | AD | ≥43 | 2 |
| Spinocerebellar ataxia 6 (MIM: 183086, SCA6) | *CACNA1A* (MIM: 601011) | CAG | AD | ≥20 | 2 |
| Spinocerebellar ataxia 8 (MIM: 608768, SCA8) | *ATXN8OS* (MIM: 603680) / *ATXN8* (MIM: 613289) | CTG/CAG | AD | ≥80 | 1 |

AD: autosomal dominant. XLR: X-linked recessive.

### Supplemental Table S7A.  Clinically significant cardiac structure/function findings

| **CHAMBER SIZES** | **NUMBER OF INDIVIDUALS** |
|---|---|
| Left atrium enlargement | 14 (moderate); 4 (severe) |
| Right atrium enlargement | 3 (moderate); 2 (severe) |
| **HYPERTROPHY** | **NUMBER OF INDIVIDUALS** |
| Mild LV hypertrophy | 119 |
| Moderate or severe LV hypertrophy | 2 (moderate) |
| Asymmetric septal hypertrophy | 1 (severe) |
| LV non-compaction cardiomyopathy | 1 |
| **RIGHT VENTRICULAR FUNCTION** | **NUMBER OF INDIVIDUALS** |
| Right ventricular systolic dysfunction | 1 (mild) |
| Pulmonary Hypertension | 12 (mild) |
| **LEFT VENTRICULAR FUNCTION** | **NUMBER OF INDIVIDUALS** |
| Left ventricular systolic dysfunction | 2 (moderate) |
| Left ventricular diastolic dysfunction | 4 (stage II) |
| Wall-motion abnormalities | 1 |
| **VALVES** | **NUMBER OF INDIVIDUALS** |
| Bicuspid aortic valve | 8 |
| Mitral valve prolapse | 10 |
| Aortic valve regurgitation | 32 (mild) |
| Tricuspid, mitral or pulmonary valve regurgitation | 3 (mitral); 2 (tricuspid); 1 (pulmonary) |
| Aortic or mitral stenosis | 6 (mild); 1 (moderate); 1 (severe) |
| **MASS OR THROMBUS** | **NUMBER OF INDIVIDUALS** |
| Abnormal mass | 2 |
| **ATRIAL OR VENTRICULAR SEPTAL DEFECT** | **NUMBER OF INDIVIDUALS** |
| Atrial septum aneurysm | 6 |
| Ventricular septal defect | 1 |
| **AORTA** | **NUMBER OF INDIVIDUALS** |
| Aortic root dilation | 18 |

### Supplemental Table S7B. Clinically Significant Findings of Arrhythmia and Conduction Disorders

| **Conduction disorders** | **Number of individuals** |
|---|---|
| Prolonged QT | 44 |
| Right Bundle Branch Block | 17 |
| Left Bundle Branch Block | 3 |
| Left anterior fascicular block | 6 |
| Bifasicular block | 3 |
| Nonspecific intraventricular conduction block | 2 |
| **Arrhythmia** | **Number of individuals** |
| Extended SVT | 45 |
| Atrial fibrillation or Atrial flutter | 15 |
| AV block (Mobitz 2nd degree type II or higher | 8 |
| Frequent Premature Atrial Contractions | 3 |
| Frequent Premature Ventricular Contractions | 2 |

SVT: Supraventricular tachycardia, AV: Atrioventricular

For atrial fibrillation or atrial flutter, eight participants (n=15) were new diagnoses.

### Supplemental Table S8. Coronary Artery Calcium Scores and Participant Characteristics

**Histogram of coronary artery calcium (CAC) score in cohort**

| CAC score | Number of individuals |
|---|---|
| **0** | 359 |
| **1 to 99** | 176 |
| **100 to 299** | 55 |
| **300 to 399** | 22 |
| **400 +** | 59 |

For participants <45 years old (n=159), 15.7% had a CAC score >1, and one participant had visible aortic root calcification. For participants with a CAC score <100 (n=512), eight individuals had a high percentile (>=90) based on their demographics (age, gender and race/ethnicity).

**Characteristics of participants with CAC score > 300**

| **High Blood Cholesterol** | Number of individuals |
|---|---|
| PMH of Hyperlipidemia or Dyslipidemia | 66 |
| Total Cholesterol (Borderline High, High) | 37 (26, 11) |
| LDL (Borderline High, High, Very High) | 23 (19, 4, 0) |
| HDL (Low) | 13 |
| Triglycerides (Borderline High, High, Very High) | 26 (13, 13, 0) |
| **Diabetes** | |
| PMH of Prediabetes or Diabetes | 17 (4, 13) |
| Glucose (Prediabetes, Diabetes) | 51 (39, 12) |
| A1C (Prediabetes, Diabetes) | 36 (28, 8) |
| **Chronic Kidney Disease** | |
| PMH of Chronic Kidney Disease | 3 |
| eGFR (Moderate, Severe) | 4 (4, 0) |
| **Hypertension** | |
| PMH of Hypertension | 33 |
| **Numbers of individuals without above criteria** | 2 |
| **Total (CT score >300)** | 96 |

PMH: Personal medical history

**Supplemental Table S9: Heatmap Representation of Blood Clinical Tests.**

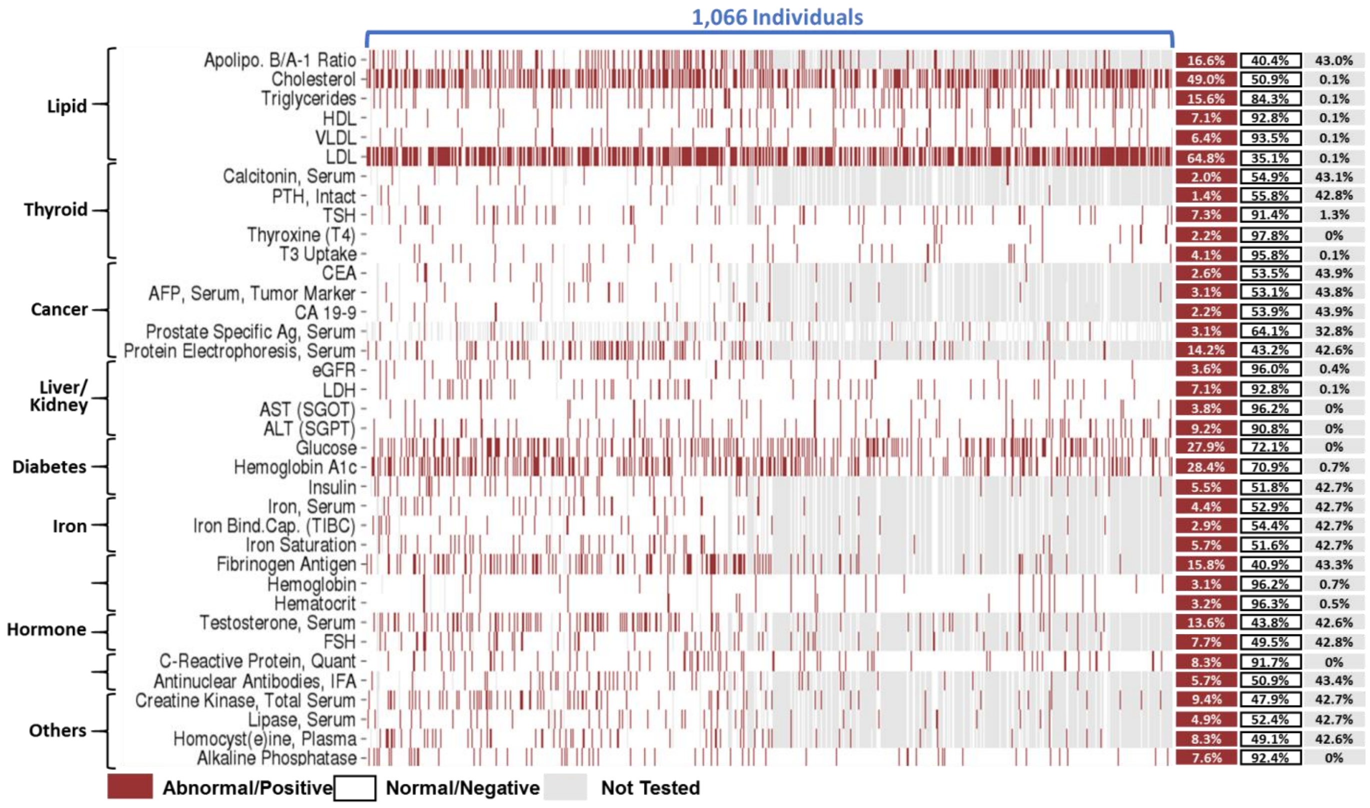


Selected test results and their grouping are shown on the left. Percentages of individuals in each finding category are color coded and listed on the right. Apolipo. B/A-1 Ratio: Apolipoprotein B/A1 ratio. HDL: high-density lipoprotein. VLDL: very low-density lipoprotein. LDL: low-density lipoprotein. PTH: parathyroid hormone. TSH: thyroid-stimulating hormone. T3: triiodothyronine. CEA: carcinoembryonic antigen. AFP: alpha-fetoprotein. Ag: antigen. eGFR: estimated glomerular filtration rate. LDH: lactate dehydrogenase. AST (SGOT): aspartate aminotransferase (serum glutamic-oxaloacetic transaminase). ALT (SGPT): alanine aminotransferase (serum glutamic pyruvic transaminase). Iron Bind.Cap. (TIBC): total iron binding capacity. FSH: follicle-stimulating hormone. IFA: indirect immunofluorescence assay.
